# Bioadhesive Hydrogel-Coupled and Miniaturized Ultrasound Transducer System for Long-Term, Wearable Neuromodulation

**DOI:** 10.1101/2024.07.17.603650

**Authors:** Kai Wing Kevin Tang, Jinmo Jeong, Ju-Chun Hsieh, Mengmeng Yao, Hong Ding, Wenliang Wang, Xiangping Liu, Ilya Pyatnitskiy, Weilong He, William D. Moscoso-Barrera, Anakaren Romero Lozano, Brinkley Artman, Heeyong Huh, Preston S. Wilson, Huiliang Wang

**Author notes:** Corresponding Author: Dr. Huiliang (Evan) Wang, Department of Biomedical Engineering, Cockrell School of Engineering, The University of Texas at Austin, Austin, Texas 787112, United States. These authors contributed equally to this work (Kai Wing Kevin Tang, Jinmo Jeong, Ju-Chun Hsieh).

## Abstract

Transcranial focused ultrasound has become a promising non-invasive approach for neuromodulation applications, particularly for neurodegenerative diseases and psychiatric illnesses. However, its implementation in wearable neuromodulation has thus far been limited due to the devices’ large size, which needs external supporting systems for the neuromodulation process. Furthermore, the need for ultrasound gel for acoustic coupling between the device and skin limits the viability for long-term use, due to its inherent susceptibility to dehydration and lack of adhesiveness to form a stable interface. Here, we report a wearable miniaturized ultrasound device with size comparable to standard EEG/ECG electrodes integrated with bioadhesive hydrogel to achieve efficient acoustic intensity upon ultrasound stimulation for long-term, wearable primary somatosensory cortical stimulation. Specifically, air-cavity Fresnel lens (ACFAL) based self-focusing acoustic transducer (SFAT) was fabricated using a lithography-free microfabrication process. Our transducer was able to achieve an acoustic intensity of up to 30.7 W/cm^2^ (1.92 MPa) in free-field with a focal depth of 10 mm. Bioadhesive hydrogel was developed to address the need for long-term stability of acoustic couplant for ultrasound application. The hydrogel demonstrated less than 13% attenuation in acoustic intensity and stable adhesion force of 0.961 N/cm over 35 days. Leveraging our bioadhesive hydrogel-integrated wearable ultrasound transducer, we were able to suppress somatosensory evoked potentials elicited by median nerve stimulation via functional electrical stimulation over 28 days, demonstrating the efficacy of our transducer for long-term, wearable neuromodulation in the brain.

## Introduction

The increase in brain diseases amongst the general population has motivated significant research in therapeutic treatment approaches. With 1 million people in the US diagnosed with Parkinson’s disease and a projected increase of 78% annually, the socioeconomic burden on individuals, families, and the healthcare system is significant^1,2^. Deep brain stimulation (DBS) has been clinically approved in treating Parkinson’s disease^3–7^, essential tremor^8–12^, epilepsy^13–16^, dystonia^17–19^ and obsessive-compulsive disorder^4,20^. Despite being a very effective method, it requires invasive implanted electrodes with complications involving hematoma, lead fractures, and glial response rendering electrodes ineffective^21,22^. Alternatively, non-invasive brain stimulation devices provide a unique opportunity for novel treatments for a multitude of psychiatric, mental, and neurodegenerative diseases in a substantial number of patients as a non-invasive intervention. Transcranial magnetic stimulation (TMS) are currently effective and clinically approved treatment methods for mental health disorders such as obsessive-compulsive disorders and depression^23^. It has also shown promising improvements in sleep disorders, Parkinson’s^24^, Alzheimer’s^25^ and potentially several other neuropsychiatric disorders^26^. However, TMS stimulates large brain areas due to its low spatial resolution, making it difficult to achieve the most effective treatment without causing adverse off-targeting effects^27–30^. Since TMS generally requires a 3∼6 weeks treatment period and DBS require continuous stimulation upon implantation, there is a strong need for non-invasive and high-spatial resolution neuromodulation approach with long-term wearability^6,25,31–33^.

Transcranial focused ultrasound (tFUS) provides an alternative non-invasive strategy for highly precise targeting of subcortical and deep brain stimulation with high spatial-temporal resolution^34^. It has shown improvement in neurological diseases such as tremor associated with Parkinson’s disease^35,36^, cognitive and memory impairments in Alzheimer’s disease^37–41^, epilepsy^42–44^, and chronic mental health disorders^45^. However, the current tFUS systems are typically bulky and are not in wearable format for long-term neuromodulation. To develop an effective wearable ultrasound neuromodulation system requires: 1) miniaturized transducer with effective acoustic intensity and focality for tFUS^46^, 2) stable fixation to the skin during neuromodulation^47^, and 3) acoustic impedance match between the ultrasound transducer and tissue^48^. Currently, the commercial ultrasound gel has been commonly used as a medium to match the acoustic impedance during ultrasound stimulation by eliminating air gaps in promoting the efficiency of ultrasound transmission^49^. Yet, its limitations of non-adhesive properties for ultrasound transducer fixation to the skin and dehydration susceptibility prevents long-term use in continuous neuromodulation treatment of brain disorders over several weeks^50^. Current approaches in developing acoustic hydrogel for wearable ultrasound imaging application has shown to be effective for ultrasound applications, however its efficacy diminishes drastically after 72 hours^51,52^. Therefore, a wearable ultrasound device integrated with an acoustically compatible medium that provides robust device-to-skin adhesion for long-term application is desired.

In this work, we developed a strategy to address the current limitations of ultrasound devices to enable long-term cortical neuromodulation. Specifically, we have developed self-focusing acoustic transducers (SFAT) that leverages geometrical patterning of acoustic lens in altering wave propagation to achieve acoustic focusing through the use of air-cavity fresnel acoustic lens (ACFAL), allowing an increase in acoustic intensity of the focal depth limit originally constrained by the geometrical diameter of the transducer (**Fig. 1a-c**) ^53^. In addition, we have designed bioadhesive hydrogel, consisting of 2-acrylamido-2-methyl-1-propanesulfonic acid (AMPS) and glycerol, to have high water absorption and rehydration properties over a month and strong adhesion to the skin (**Fig. 1d**). Through integration of bioadhesive hydrogel to our SFAT transducer, our **Min**iaturized and B**i**oadhesive-coupled **Ul**trasound **Tra**nsducer (MiniUlTra) weighs 8.5 grams and can be easily attached to the target skin for an extended period, allowing ease of use for long-term applications (**Fig. 1e-h**). We evaluated the efficacy of MiniUlTra in its effectiveness in suppressing somatosensory evoked potential elicited by median nerve stimulation via functional electrical stimulation over 28 days, demonstrating the efficacy of MiniUlTra in long-term cortical neuromodulation as a wearable ultrasound device.

**Figure 1.**
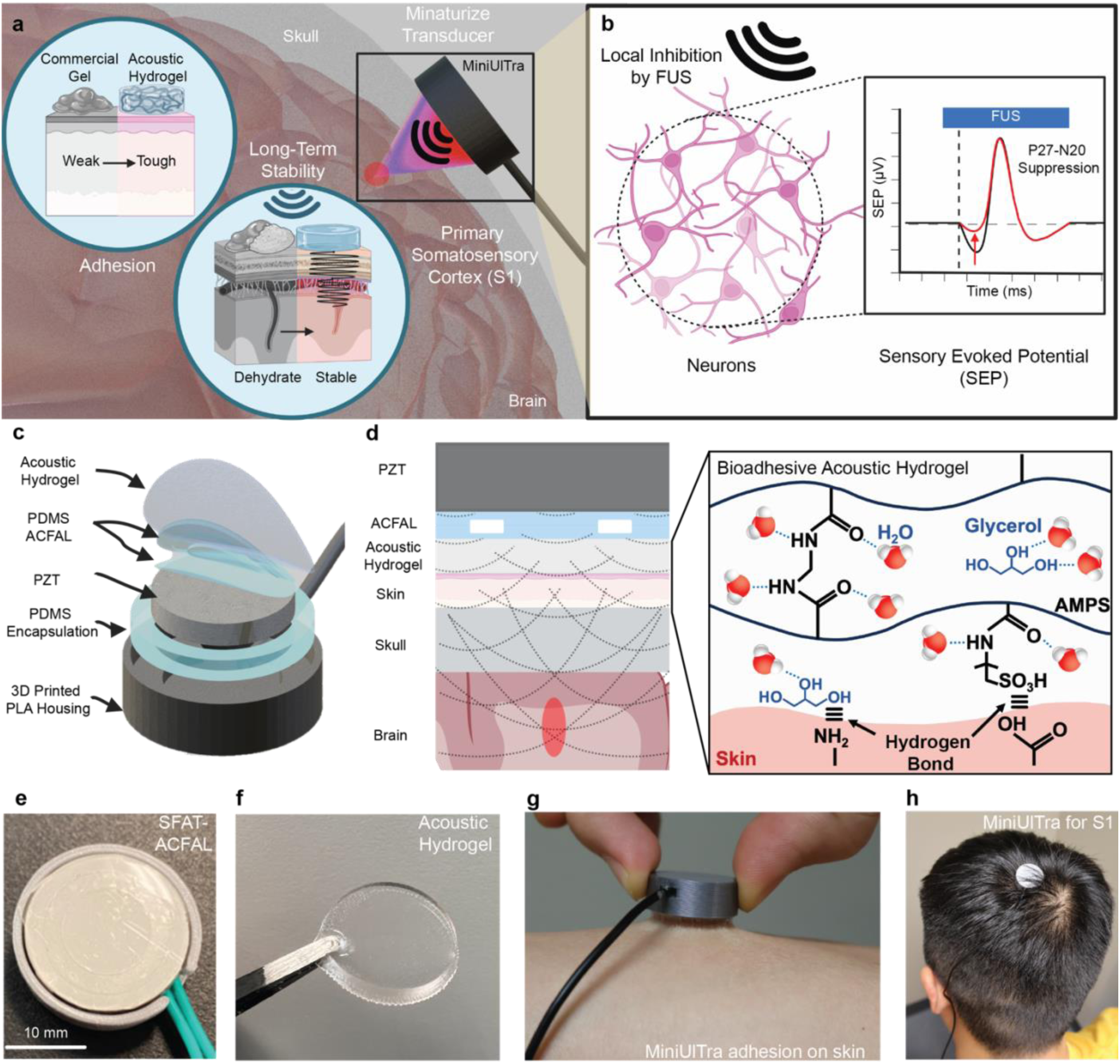
Miniaturized and Bioadhesive-Coupled Ultrasound Transducer (MiniUlTra). **a)** Illustration of MiniUlTra that continuously adheres to the scalp targeting the primary somatosensory cortex (S1) with high adhesion force, low acoustic attenuation and miniaturized transducer. **b)** Mechanism of suppression of P27-N20 complex in somatosensory evoked potential (SEP) through focused ultrasound stimulation locally at the S1. **c)** Schematic of layered structure of MiniUlTra that assembles the piezoelectric with PDMS-based ACFAL and bioadhesive hydrogel integrated into a compact 3D-printed housing. **d)** Side-view of layered schematic including chemical structure of bioadhesive hydrogel and its’ adhesion mechanisms **e-f)** Optical images of design and fabricated SFAT-ACFAL and bioadhesive hydrogel**. g)** Adhesion of MiniUlTra on skin. **h)** Demonstration of MiniUlTra on the scalp for S1 targeted neuromodulation.

## Results

### Development and characterization of miniaturized ultrasound transducer

Acoustic frequencies used for ultrasound stimulation with ideal transcranial transmission and brain absorption has been reported to be within the 500 kHz to 750 kHz range^54,55^, where clinical demonstration of using ultrasound stimulation at the 650 kHz frequency in humans has shown effective suppression evoked potential and enhanced sensory functions^56^. Thus, we developed a custom miniaturized 650 kHz ultrasound transducer (similar size to standard EEG electrode leads, OD = 18 mm) with microfabrication techniques to create a self-focusing acoustic transducer (SFAT) using air cavity Fresnel acoustic lens (ACFAL) coupled with a novel highly adhesive and conformable bioadhesive hydrogel for long-term applications. Characterization of acoustic pressure fields emitted from our SFAT-ACFAL was done using a calibrated hydrophone (**Fig. 2a**) on a motorized 3-axis system submerged in a degassed distilled water bath. Comparison of acoustic field distribution with and without the ACFAL formed by PDMS (Pristine PZT/SFAT-ACFAL) showed less scattering and higher focusing on the desired focal point in the transducer with ACFAL (**Fig. 2b**). Recording of acoustic waveforms pulsed and transmitted was performed in free-field and with a macaque skull, where measurements indicate a spatial focality of 3.5 mm axially and 8 mm radially (**Fig. 2c-d**). The focal depth was measured to be at 10 mm, at the expected and designed specification. To determine the acoustic intensity and biosafety of the devices for ultrasound neuromodulation, a calibration curve was performed with our ultrasound system (Image Guided Therapy System) used to drive the SFAT-ACFAL to evaluate the linearity of acoustic intensity and pressure when driving amplitude was increased. The measurements revealed a spatial-peak pulse-average intensity (I_SPPA_) in the free field to be less than 30.7 W/cm^2^ (1.92 MPa) (**Fig. 2e**). However, free-field acoustic intensity of 23.9 W/cm^2^ has shown to be effective when transmitting through the human skull with a four-fold drop in intensity to 5.9 W/cm^2^ ^56^. Therefore, all stimulation paradigms performed in healthy volunteers using our device were performed at 23.1 W/cm^2^ (1.66MPa), to necessitate sufficient acoustic intensity threshold in suppression of sensory evoked potentials^56^. The effect of transcranial skull transmission effectively attenuates the amplitude of the acoustic pulse waveform (**Fig. 2f**) and increases the spatial resolution of the focal spot. This is suspected due to the inhomogeneity of skull and tissue interfaced between the boundary conditions resulting in time-reversible wave propagation, commonly used for imaging^57^. The shift in axial peak of the focal spot (**Fig. 2d**) was due to the curvature of the macaque skull and its difficulty in positioning between the hydrophone and device to prevent collision.

**Figure 2.**
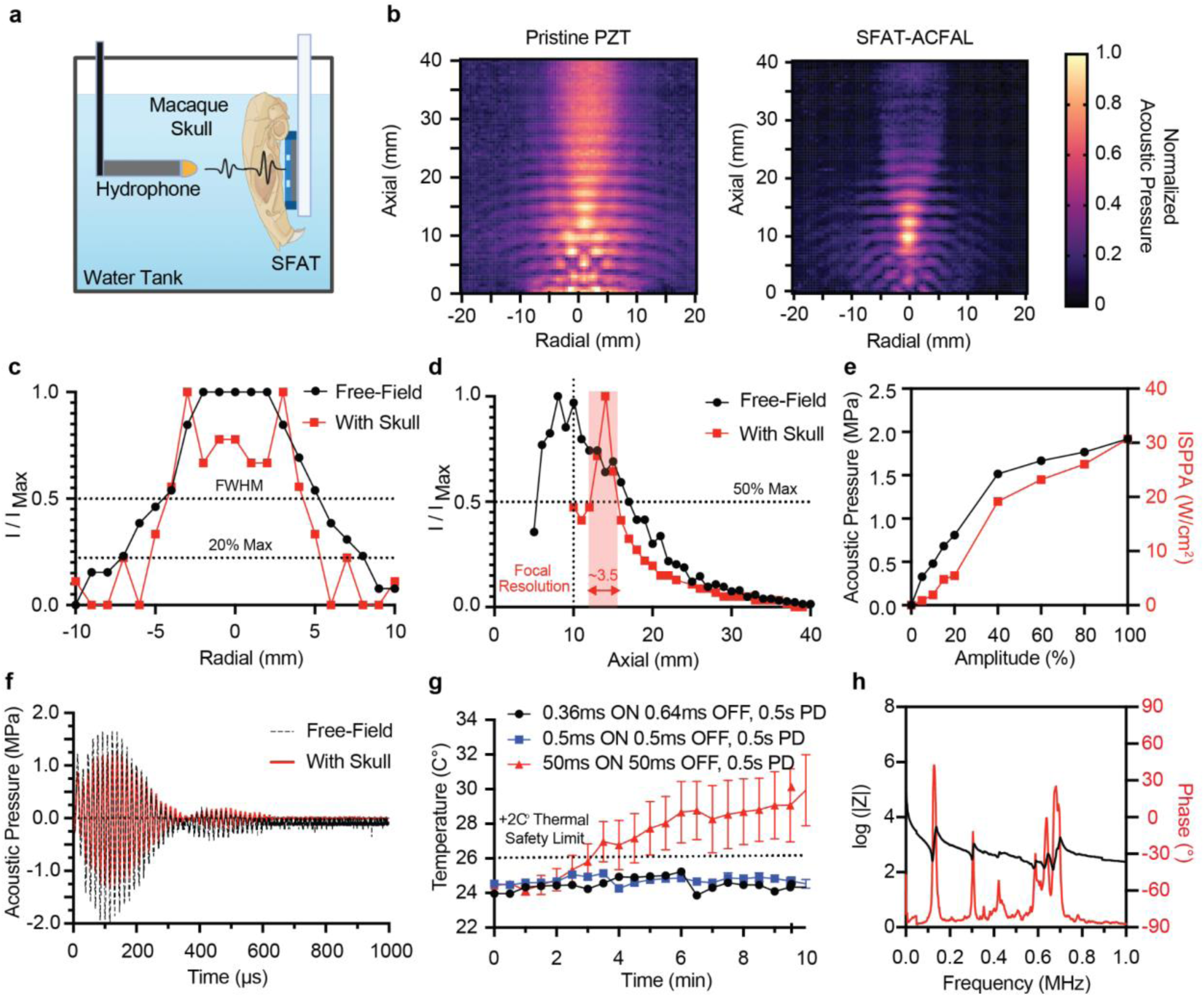
Self-Focusing Acoustic Transducer (SFAT) using Air-Cavity Fresnel Lens (ACFAL). **a)** Schematic of experimental setup for characterization of SFAT-ACFAL. **b)** Comparison of acoustic field distribution and intensity with Pristine PZT (left) and with PDMS-based ACFAL (right). **c)** Normalized radial acoustic intensity profile in free-field and with the presence of a macaque skull at focal depth 10 mm. **d)** Normalized uniaxial acoustic intensity profile in free-field and with the presence of a macaque skull. **e)** Acoustic pressure (MPa) and intensity (I_SPPA_) calibration curve measured when SFAT-ACFAL at varying driving amplitude using ultrasound generator system. **f)** Measured waveform of ultrasound pulse using stimulation paradigm of 360µs with and without macaque skull. **g)** Thermal effect of SFAT-AFAL on macaque skull measured with infrared camera on varying stimulation parameters. **h)** Electrical impedance and phase of SFAT-ACFAL.

To ensure thermal biosafety, the device during ultrasound stimulation should not exceed an increase of 2°C ^58^. We then characterized the thermal biosafety of the device by performing the stimulation paradigm used by Legon *et al.*^56^ in comparison with higher duty cycle and pulse duration through the macaque skull and monitoring the temperature of the stimulation site using an infrared camera (**Fig. 2g, Supplementary Fig. 1**). Results indicated that with 10 min of continuous stimulation, the paradigm of 360 µs ON and 640 µs OFF ^56^ had no thermal increase. Similarly, when the duty cycle was increased to 50% (500 µs ON and 500 µs OFF) there was no significant temperature increase. However, when the pulse duration increased to 50 ms ON and 50 ms OFF, drastic temperature increase was observed beyond the 5 min mark. This demonstrates the safety regime of the device for human applications. Electrical characterization of the device was performed using an impedance spectrum analyzer to determine the impedance and phase response of SFAT-ACFAL. The impedance was then used to determine harmonic frequencies where impedance is lowest for impedance matching purposes (**Fig. 2h**).

### Development and characterization of bioadhesive hydrogel

To ensure long-term neuromodulation viability, the need for an acoustic couplant to sustain stable acoustic properties over time is needed to be integrated with the SFAT-ACFAL. Specifically, it should sustain hydration, effectively transmit ultrasound, adhere conformally to the skin with a low modulus to minimize air gaps, and maintain high adhesion force over time^51,52^. The bioadhesive hydrogel used in this study includes two primary materials: 1) 2-acrylamido-2-methylpropane sulfonic acid (AMPS) and 2) glycerol (**Fig. 1d**). PolyAMPS is an ionic polymer with a hydrophilic sulfonic group resulting in it being inherently negatively charged, which allows for strong ionic interaction with water molecules^59^. Thus, it enables high water absorption rate^60–62^, allowing sustained hydrated state through absorption of ambient moisture^63^. In addition to its high water content and retention, PolyAMPS provides modulus similar to that of biological tissues^64^, and is suitable as a long-term substitute of commercially available ultrasound gel that tends to dehydrate within hours. Furthermore, the addition of glycerol containing hydroxyl groups, which forms hydrogen bonds with water molecules, provides water retention capacity and enhanced adhesion to the skin by offering a hydrating effect on the stratum corneum ^60,65,66^.

We characterized the acoustic attenuation rate of our bioadhesive hydrogel at different thickness in comparison to commercial gel. As a result, we observe that the ultrasound power attenuation of the bioadhesive hydrogel is comparable to that of commercial gel at thicknesses of 0.5 mm and 1 mm **(Fig. 3a)**. With ultrasound as a mechanical acoustic wave, the mismatch in impedance when propagated between mediums with varying acoustic impedances inevitably leads to partial transmission and reflection at the boundary layers. By minimizing the mismatch in impedance, reduction of reflected and maximizing transmitted waves provides higher acoustic intensity deposition to the target site. Therefore, the need for minimizing the impedance mismatch between our device and the skin using our bioadhesive hydrogel is necessary^51^. As acoustic speed in a material is directly related to its elastic properties and density, the acoustic impedance could be derived directly from the acoustic speed and density^67,68^. We experimentally characterize and measure the acoustic speed of our hydrogel. Firstly, with the acoustic speed of water being 1500 m/s and the distance between PZT and hydrophone positioned 20 mm apart in the water tank, the time required for an acoustic pulse (time-of-flight, ToF) to travel from the transducer to the hydrogel theoretically will be 13.3 µs in free-field. Upon measurement of ToF with the hydrogel, comparison of to the free-field measurement allows us to determine the time difference and the acoustic speed through the hydrogel **(Fig. 3b)**. The measured density of the hydrogel was 1166.7 kg/m^3^. Overall, the acoustic impedance of the hydrogel yielded 2.13 ± 0.11 MRayl and 2.17 ± 0.13 MRayl, with estimated acoustic speed of 1816 ± 76.36 m/s and 1864 ± 113.7 m/s on day 0 and day 7 respectively (**Fig. 3c, Supplementary Fig. 2**). Overall, the acoustic impedance of the bioadhesive hydrogel remained stable across 7 days, with an average hydrogel impedance of 2.17 MRayl. Compared to the acoustic impedance of 1.99 MRayl for skin^52^, our bioadhesive hydrogel exhibits a much more similar impedance to human skin compared to commercial ultrasound gels such as Konix Sterile Gel and Aquasonic 100, indicating minimum acoustic loss of our hydrogel in addition to its’ long-term stability.

**Figure 3.**
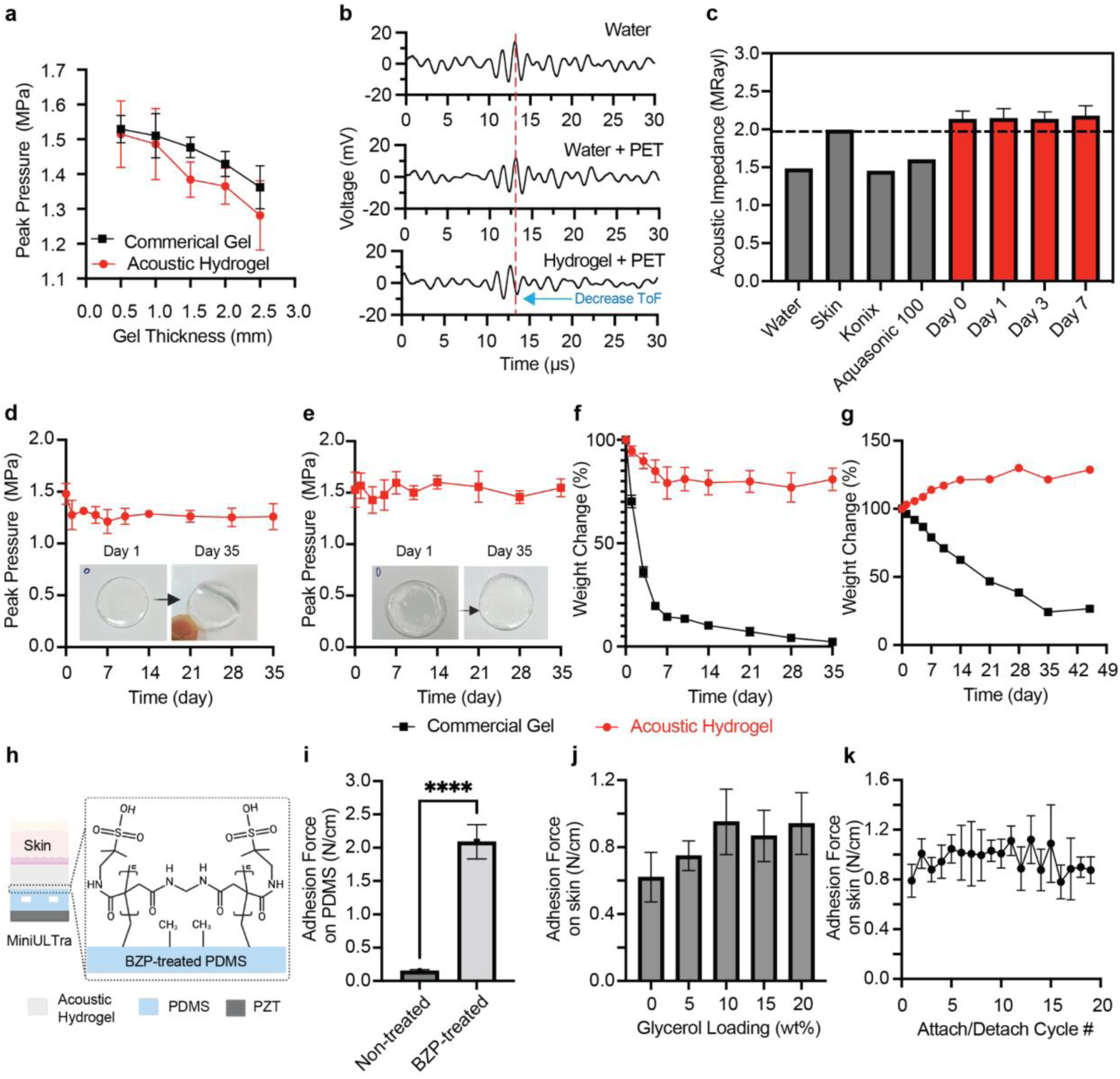
Bioadhesive hydrogel. **a)** Comparison of ultrasound intensity decreases according to thickness changes in bioadhesive hydrogel and commercial gel (Aquasonic 100, Parker), (n = 6 for each thickness). **b)** Comparison of acoustic time-of-flight (ToF) for ultrasound transmission through water, PET, and hydrogel. **c)** Acoustic impedance of the hydrogel for 7 days (n = 4 each day) compared to commercial gel, water, and human skin. **d)** Peak-pressure attenuation of bioadhesive hydrogel under 30% humidity and 22°C over 35 days. (n = 4). **e)** Peak-pressure attenuation of bioadhesive hydrogel under 75% humidity and 22°C over 35 days. (n= 4). **f)** Weight change of the bioadhesive hydrogel and commercial gel in the room environment (humidity: ∼30 %, temperature: ∼22 °C, n = 4). **g)** Weight change of the bioadhesive hydrogel and commercial gel with high humidity (humidity: ∼75 %, temperature: ∼22°C, n = 4). **h)** Chemical structure of the bioadhesive hydrogel integrated ACFAL by grafting the bioadhesive hydrogel to benzophenone (BZP) treated PDMS. **i)** Improvement of adhesion force with BZP-treated PDMS (n = 4). **j)** Adhesion force of the bioadhesive hydrogel according to glycerol loading change (n = 5). **(k)** Adhesion force of the 20-cycle attachment/detachment test of the bioadhesive hydrogel on skin (n=4).

Further investigation of the long-term stability of the hydrogel in acoustic attenuation was performed under low and high humidity conditions. Low humidity of 30% mimicked a typical indoor room environment and 75% reflects high humidity outdoor conditions, which was emulated by a sealed humidity controlled container storage where the hydrogel was stored^69,70^. Our hydrogel had an attenuation of up to 13% and less than 2% when stored in humidity conditions of 30% and 75% respectively over 35 days (**Fig. 3d-e**). Additionally, weight loss test was also performed for both our hydrogel and a commercial gel stored in these two humidity conditions. Under low humidity (RH 30%), our hydrogel exhibited a slow dehydration rate, retaining 76% of its weight and remained stable post 7 days. Conversely, the weight of the commercial gel significantly decreased with only 14% weight retention on day 7 (**Fig. 3f**). Under high humidity (RH 75%), our hydrogel had a significant and consistent increase in weight of approximately 120% after two weeks (**Fig. 3g**). Additional tests under room temperature and humidity conditions exposure were also performed. Our hydrogel exhibits a slow dehydration rate, retaining ∼65% of its volume after 24 hours. In contrast, the commercial gel undergoes rapid dehydration, retaining only ∼63% volume after 3 hours and approximately 10% weight after 24 hours. After 24 hours, the commercial gel is nearly fully dehydrated, whereas the hydrogel remains stable (**Supplementary Fig. 3 and 4**). This suggests the potential of the self-recoverable and self-rehydrating hydrogel properties within high humidity conditions.

To ensure long-term robustness in wearability, sufficient adhesion between SFAT-ACFAL and the skin is necessary. Additionally, strong adhesion between the hydrogel and the PDMS-based ACFAL is required aside from the interface between hydrogel and the skin to prevent detachment. The robust integration between ACFAL and hydrogel was achieved by using the photografting agent for the hydrogel. Treating PDMS with benzophenone (BZP), a type II photoinitiator, extracts hydrogen from the grafted surface of PDMS and generates radicals allowing the PDMS and hydrogel to form a polymeric bond under UV irradiation^71^ (**Fig. 3h**). As a result, the adhesion of BZP-treated PDMS to the hydrogel was 2.09 N/cm, which was 13 times higher than the adhesion of non-treated PDMS to the hydrogel (0.1513 N/cm) (**Fig. 3i**). Optimization of the adhesiveness of our hydrogel to the skin was achieved by tuning the loading of glycerol and was determined via measurement of adhesion force through 90° T-Peel test. As the glycerol loading increases, the adhesion force of the hydrogel improves and plateaus when glycerol loading exceeds 10 wt% (**Fig. 3j**). In this study, glycerol was loaded at 20 wt% to maintain high water retention properties, allowing an adhesion force of ∼0.941 N/cm, sufficient for attachment to the skin. Skin adhesion cycling was performed subsequently to determine the adhesive reusability, where adhesion force remained stable over 20 cycles with a mean adhesion force of 0.961 N/cm (**Fig. 3k**). Modulus compliance of hydrogel with skin was investigated, where the modulus of the hydrogel is ∼31.4 kPa, similar to that of skin tissues. Thus, providing minimal mechanical mismatch and demonstrating suitability of skin-device interface (**Supplementary Fig. 5**)^72^. These results indicate that the bioadhesive hydrogel could provide an alternative to long-term ultrasound applications.

### SFAT-ACFAL enables suppression of somatosensory evoked potentials at S1 targeting

Studies in SEP by median nerve (MN) stimulation have been explored and researched greatly. Well-defined characteristic morphology of EEG signals of SEP are distinguished into waveform peaks assigned by their polarity (positive P or negative N) and its corresponding post-stimulus latency (in ms)^73^. The changes in latency and amplitudes of these waveform peaks are often interpreted as dynamical alterations in neural activity because of combination from peripheral and central nervous system to external stimulus. Specifically, early SEP peaks or “short latency” SEPs occurring within 40 ms post-stimuli are of great importance as they have the least variability in response to peripheral external stimulation whereas long latency responses are susceptible to cognitive factors and higher ordered complex neural processing of the sensory pathways^73^. Thus, waveform peaks of N20, P27, N33, P50, N70, P100 and N140 were examined. In brief, each of these peaks serve as a biomarker with implications of tactile information processing. However, of most great interest corresponds to N20 (or commonly known as P27-N20 complex) has been highly known for its relevance to the sensory input of dorsal column-medial lemniscal pathway and acts as a primary evoked response in response to peripheral stimuli to the lateral portion hand area of somatosensory cortex extended posteriorly over to supramarginal gyrus^74^.

Recently, transcranial focused ultrasound stimulation at the somatosensory cortex has been shown to suppress somatosensory evoked potential (SEP) via the elicitation of sensory stimuli. The suppression in SEP effectively resulted in higher subjects’ ability to discriminate fine differences in two points through sensory perception at the epidermis of distal phalanges^56^. To demonstrate the efficacy of our SFAT-ACFAL, we applied our device in targeting the left S1 through transmission of tFUS into the cortex at the CP3 site (**Fig. 4a**). Electroencephalographic (EEG) electrodes using commercial Ag/AgCl was applied at the scalp of electrode sites CP1, C3, P3, and CP5 in the 10-20 EEG configuration as a means to study the influence of tFUS short-to-late onset evoked brain activity through understanding of changes in peak-to-peak amplitudes of SEP complexes and spectral changes in power elicited by the contralateral (right) MN stimulation with functional electrical stimulation (FES) (**Fig. 4b**).

**Figure 4.**
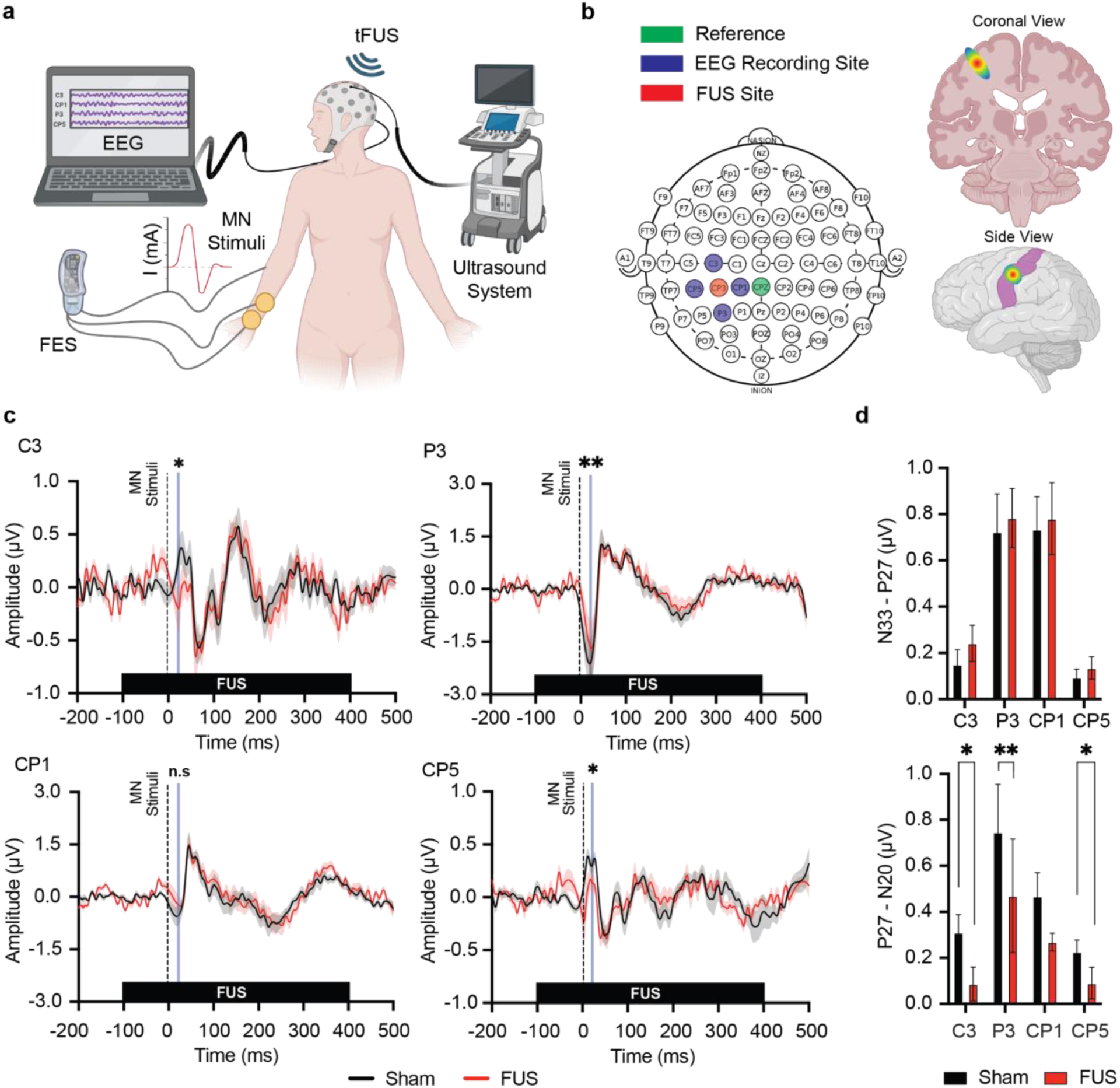
Evaluation of neuromodulation in somatosensory evoked potential using SFAT-ACFAL. **a)** Schematic representation of experimental setup. **b)** Illustration of EEG electrode and SFAT-ACFAL placement in 10-20 EEG montage with its corresponding targeting of left S1 with FUS at CP3. **c)** Grand average of epochs where median nerve (MN) stimulation occurs at t = 0 ms and FUS or sham begins at t = −100 ms (red line indicates SEP under FUS+, black line indicates SEP under sham, highlighted blue indicates where significant difference was observed P27-N20 complex in SEP). **d)** Summary of effect of FUS compared to sham in P27-N20 and N33-P27 complexes of SEP. Suppression of early onset P27-N20 complex observed across C3, P3, and CP5 in SEP by FUS (n=5 per group, Wilcoxon signed-rank test, 4 male and 1 female). All plots show mean ± s.e.m unless otherwise mentioned, *P < 0.05, **P < 0.01, and ***P < 0.001.

To determine the efficacy in SEP suppression of our miniaturized transducer, we first applied commercially available ultrasound gel coupled between the SFAT-ACFAL with the scalp at the 10-20 EEG electrode site CP3. 650 kHz tFUS beams were pulsated to the target region (n = 5) with a pulse of 360 µs ON and 640 µs OFF at a pulse repetition frequency (PRF) of 1 kHz for 500 ms. The stimulation paradigm chosen has been demonstrated experimentally in humans to suppress SEP^56^ whilst ensuring minimal thermal heating effects with our device due to the short pulse time (**Fig. 2g**). MN stimulation occurred for 200 µs at 100 ms after the beginning of tFUS transmission. Sham and tFUS treatment conditions were performed identically apart from the device being turned off in the sham group. Some subjects reported auditory chirping noises initially at the beginning of each trial produced by the device during stimulation. However, the chirping noises quickly subsided within a few seconds reported by subjects. Additionally, subjects did not report any discomfort, heating, or abnormal sensations at the site of tFUS treatment between sham and tFUS treatments.

With our device for tFUS treatment, we demonstrated a significant decrease in short latency peaks (P27-N20 complex) across electrode sites at C3 (sham, 0.289 ± 0.083 µV s.em. ; tFUS 0.086 ± 0.073 µV s.e.m), P3 (sham, 0.803 ± 0.221 µV s.em. ; tFUS 0.469 ± 0.247 µV s.e.m), and CP5 (sham, 0.222 ± 0.056 µV s.em. ; tFUS 0.089 ± 0.068 µV s.e.m) compared to the sham. Conversely, no significant reduction in short latency peaks were observed at electrode site CP1 (sham, 0.458 ± 0.109 µV s.em. ; tFUS 0.266 ± 0.038 µV s.e.m). tFUS using SFAT-ACFAL did not produce any significant changes in long-latency peaks (**Fig. 4c-d, Supplementary Table 1-4**) but late potential (>140 ms) showed general attenuation across all electrodes in late-onset SEP complexes.

Spectral decomposition of EEG signals enables understanding of spatial-temporal changes in dynamics regarding excitation and inhibition of cortex in response to information processing^75,76^. Therefore, spectral analysis was performed on the grand averaged epochs of SEP to evaluate the effects of tFUS using SFAT-ACFAL. By taking the difference between the spectral decomposition of FUS and sham, a significant short latency decrease in alpha (7-12 Hz) and beta (13-30 Hz) band power of −6 dB was observed within 100 ms of MN stimuli. Additionally, a short period of low gamma band (30-50 Hz) power decrease was observed around 100 ms post MN stimuli (**Supplementary Fig. 6**).

### Long-term wearability and neuromodulation of MiniUlTra

The ability for our device to stimulate the S1 region long-term was tested within the same experimental protocols that target the characteristic pattern of SEP with MN stimuli. Particularly, SEP suppression via tFUS tests were conducted in 3 sessions (Day 1, 7, and 28) throughout 28 days (**Fig. 5a**) to investigate the efficacy of neuromodulation using our MiniUlTra device (bioadhesive hydrogel incorporated) on healthy volunteers (n = 4). The length of the experimental protocol was chosen to investigate the extreme longitudinal conditions of our MiniUlTra device over a month period, where our bioadhesive hydrogel remains stable compared to commercial gel over 28 days (**Fig. 5b**). When MiniUlTra was used on day 1, significant suppression of SEP was observed compared to the sham group across all electrode channels. Furthermore, there was no significant difference between the treatment group when our bioadhesive hydrogel was used compared to commercial gel, validating the acoustic characteristics of the bioadhesive hydrogel in ultrasound transmission has similar performance to commercially available ultrasound gel (**Fig. 5d**). Additional sessions were performed on day 7 and day 28, which also showed significant suppression against the sham condition across all channels except for CP1 on day 7. We also observe a general decrease in the P27-N20 complex over time. Overall, the SEP amplitude across the epoch was observed with clear decreases in short (P27-N20 complex) and long latency (>70 ms) biomarkers (**Fig. 5c**). However, long latency biomarkers are more complex in its relation with median nerve stimuli due to its association with indirect somatosensory pathways involving cognitive and motor processes^77^. Hence, the P27-N20 complex was focused due to its prominent and well established association to contralateral stimuli at the S1 region^78,79^. Results indicated significant reduction in amplitude at the corresponding P27-N20 complexes across all electrode channels over 28 days (**Fig. 5d**), demonstrating robustness in ultrasound neuromodulation over long time stimulation with our MiniUlTra.

**Figure 5.**
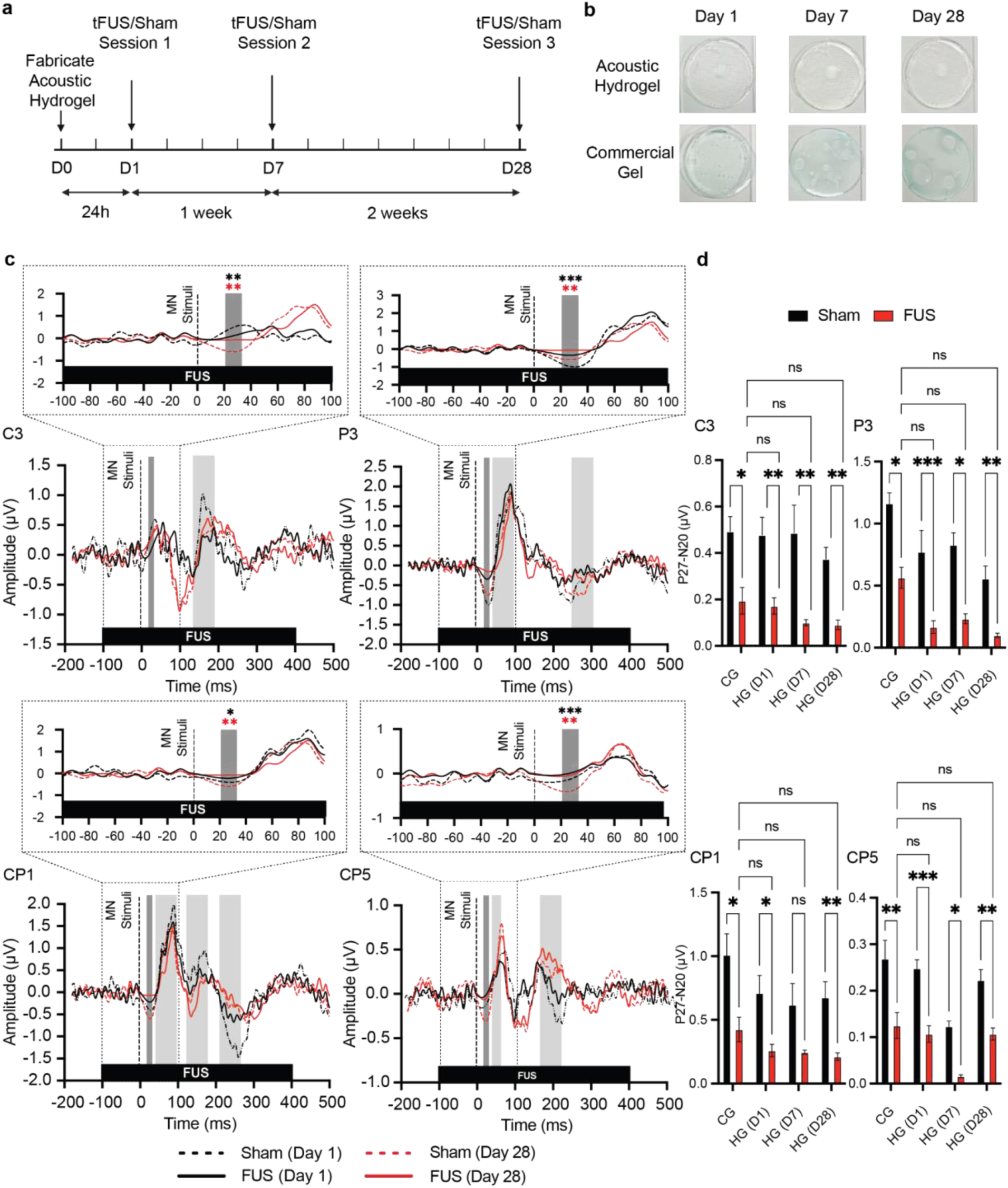
Long-term suppression of the P27-N20 complex in somatosensory evoked-potential (SEP) using MiniUlTra. **a)** Long-term experimental protocol for evaluating efficacy of hydrogel. The hydrogel was fabricated a day before the first session (D0). Three sessions per subject, each consisting of 10 trials of 3 minutes, each trial consisting of 120 epochs (tFUS/Sham) on day 1 (D0), 7 (D7) and 28 (D28). Each subject had their personal hydrogel with the device, which was stored in room temperature and ∼30% humidity. **b)** Optical image of prepared hydrogel compared to commercial ultrasound gel with the corresponding sessions. **c)** Grand average epochs comparing effects of sham and FUS with median nerve (MN) stimulation on SEP across days 1, 7 and 28 using hydrogel. MN stimulation occurs at t = 0 ms and FUS or sham begins at t = −100 ms (black solid line indicates SEP under FUS+ on day 1; black dashed line indicates SEP under Sham on day 1; red solid line indicates SEP under FUS+ on day 28; red dashed line indicates SEP under Sham on day 28; highlighted dark gray indicates where significant difference was observed P27-N20 complex in SEP; highlighted light gray indicates where observable differences in long latency complexes). **d)** Suppression of early onset P27-N20 complex observed across C3, CP1, and CP5 in SEP by FUS shown within each group. No significant difference was observed across FUS groups in bioadhesive hydrogel (HG) when compared to the FUS group using commercial gel (CG). Significant decrease in P27-N20 complexes was observed when comparing within each day of the hydrogel. (n = 4 per group, Two-Way Anova, 4 male). All plots show mean ± s.e.m unless otherwise mentioned, *P < 0.05, **P < 0.01, and ***P < 0.001.

## Discussion

We have demonstrated a newly developed bioadhesive hydrogel coupled and miniaturized wearable ultrasound transducer that offers long-term brain neuromodulation capability without the need of handheld operators and fixtures. The device utilizes an alternative simplified microfabrication approach without the need of standard lithography techniques for SFAT-ACFAL patterning to achieve higher focality, acoustic intensity, and miniaturization. Additionally, our development of a novel hydrogel provides mechanical compliance, bioadhesion and stable acoustic coupling between our device and skin interface. For the first time, our hydrogel has shown acoustic and adhesive stability for more than a month compared to current state-of-the-art acoustic hydrogels stability of 72 hours. By integrating the two components, our device MiniUlTra can be used to perform noninvasive focused ultrasound stimulation delivered into the cortical region over 28 days with robust performance and clinical applications. Biosafety of the device was demonstrated to achieve spatial pulsed averaged intensity and acoustic pressure within the safety limits suggested by FDA guidelines and literature. Thus, our system provides a promising platform for non-invasive long-term wearable ultrasound applications.

In conclusion, wearable ultrasound stimulation devices hold significant promise for the long-term treatment of chronic diseases like Parkinson’s disease, essential tremor, epilepsy and depression. These devices offer non-invasive, spatiotemporal targeted modulation of neural activity, potentially improving disease symptoms without the drawbacks of medications or surgery. Their non-invasive nature and wearability also suggest the potential for home-based therapy, although continued research is essential to optimize treatment protocols and ensure long-term safety and efficacy across diverse patient populations.

## Methods

### Fabrication of SFAT-ACFAL

Geometric shape and radius of the ACFAL was determined first by selection of 10 mm focal depth according to the equations governed by Fresnel lens^80^, which was then implemented into finite element analysis software for simulation (COMSOL Multiphysics 6.0, COMSOL Inc.). Optimization of PDMS and air-cavity thickness was performed with reference to previous feasibility of microfabrication (**Supplementary Fig. 7**)

Mold glass substrates were initially patterned by first laminating 36 um thick copper tape (1125, 3M) onto adhesive interlayer (Ultra 582U, TransferRite), which was then laminated onto an adhesive backing layer (GXF341, DigiClear Plus). The laminated copper tape was then negatively patterned using laser etching (LPKF, U4 Laser) and transferred printed onto the glass substrate (**Supplementary Fig. 8i**).

Patterned mold glass substrates were cleaned and prepared by first submerging into a beaker filled with acetone and sonicated to remove particulates for 5 min. Substrates were then removed, rinsed with distilled water and submerged in methanol for 5 min of sonication. The substrates were then rinsed with distilled water before blow dried with purified nitrogen gas. Substrate was spin-coated with a sacrificial layer (Omnicoat, Kayaku Advanced Materials) for 30s at 1000 RPM and 3 min of planarization before soft-baking at 200°C on a hotplate. The parameters were determined empirically through patterning and measurement of thickness using profilometer (**Supplementary Fig. 8**) Subsequently, substrates were then spin-coated with 5 ml of PDMS (Sylgard 184); prepared by mixing 1:10 of curing agent with base elastomer and desiccated for 1 hour at 500 RPM to achieve ∼200 µm thickness and cured on a hotplate at 90°C for 35 mins. Substrates were then placed in acetone filled beakers and sonicated for 5 min each to release the patterned PDMS mold (**Supplementary Fig. 8ii**). Using tweezers, the patterned PDMS layer was carefully removed and placed onto a temporary glass substrate, which was then trimmed with medical scalpel. Similarly, the PZT (DL-47, Del Piezo) was subjected to the same substrate cleaning process mentioned previously. 2 ml of prepared PDMS was spin-coated onto the surface of PZT at 2000 RPM for 30s to achieve a thickness of 40µm; cured at 90 °C for 30 min (**Supplementary Fig. 8iii**).

Next, the released patterned PDMS layer and coated-PDMS PZT was treated with Reactive Ion Etching (RIE) O_2_ plasma treatment for 25 s (30W @ 30% O_2_, 30 SCCM) to remove organic hydrocarbons on the surface and create silanol (SiOH) functional groups, effectively increasing the wettability and rendering surface more hydrophilic^81^. The patterned PDMS layer was then reversely bonded onto the coated-PDMS PZT by attachment and applying 1 kg weight simultaneously on a 120 °C hotplate for 5 min (**Supplementary Fig. 8iv**).

### Acoustic Field Mapping of SFAT-ACFAL

#### Mapping

The SFAT-ACFAL device was mounted on a submersible stand in a degassed distilled glass water tank. Acoustic intensity and waveform were measured using a calibrated capsule hydrophone (HGL-0200, Onda) mounted on a three-axis stage system, which was connected to an oscilloscope (SDS 1204-XE, Siglent) via a signal preamplifier (AG-2010, Onda) interfaced to a custom MATLAB program for automated 3D scanning and signal processing (**Supplementary Fig. 9**). The device was controlled and actuated by a commercially available ultrasound system (BBBoq, Image Guided Therapy Systems). Acoustic field scans without macaque skulls were first performed at 500 µm increments (0 - 40 mm from transducer in a 40 mm x 40 mm grid workspace). Focal depth and spatial peak locations were obtained from the acquired acoustic field scans axially and radially. Subsequently, the macaque skull (3-mm thick macaque cortical bone, rehydrated for 24 h in phosphate buffer solution) was inserted in between the transducer and hydrophone using the same scan procedures. Due to the curvature and inhomogeneous geometry, acoustic field scans were performed at 500 µm increments (∼10 - 40 mm from the transducer in a 40 mm x 40 mm grid workspace) to avoid collision between transducer, skull, and hydrophone.

#### tFUS Waveform

Generation of tFUS profile from SFAT-ACFAL was performed using a 40-W high-voltage biphasic ultrasound function generator system (BBBoq, Image Guided Therapy System) controlled and pulsed by an external Arduino trigger. Briefly, the function generator was set to deliver individual pulses at 360 µs ON and 640 µs OFF with center frequency of 650 kHz (**Fig. 2e**). The Arduino was then programmed to trigger the function generator at a pulse repetition frequency (PRF) of 1 kHz and pulse duration of 500 ms ON and 500 ms OFF.

#### Electrical Characteristics

SFAT-ACFAL was connected to an impedance spectrum analyzer (SP300, BioLogic) using a two-electrode connection configuration. Impedance of the device was measured from 0-1MHz to validate resonant frequencies. Fundamental harmonics and phases were identified in addition to the desired 650kHz (**Figure 2h)**.

#### Thermal Heating

SFAT-ACFAL was placed facing upwards on a 3D-printed mounted stand, where the superficial side of the macaque skull was placed in contact with the transducer using ultrasound coupling gel (Aquasonic 100, Parker). Three stimulation paradigms with varying duty cycle and pulse duration were used (360 µs ON/640 µs OFF, 500 µs ON/500 µs OFF, 50ms ON/50ms OFF) for 10 mins to compare and observe the thermal heating effects from tFUS (**Fig. 2f**). An infrared camera (One Edge, FLIR) was used to record three points in a triangular configuration surrounding the targeting area on the inferior side of the macaque skull (**Supplementary Fig. 1**).

### Synthesis and integration of bioadhesive hydrogel to SFAT

#### Materials and fabrication of bioadhesive hydrogel

The preparation of the bioadhesive hydrogel started with mixing the hydrogel solution. First, AMPS (Sigma-Aldrich) was dissolved in deionized (DI) water at a 1:1 ratio using a vortex mixer for 30s. Subsequently, glycerol (Alfa Aesar) with 20 wt% was added to the AMPS/DI water mixture using a vortex mixer for 30s. N, N’-Methylenebis(acrylamide) (MBAA crosslinker, Sigma-Aldrich) with∼0.16 wt% was then added and mixed for 60s. Irgacure 2959 (2-Hydroxy-4’-(2-hydroxyethoxy)-2-methylpropiophenone 98%, Sigma-Aldrich) with ∼0.59 wt%, serving as the photoinitiator, was mixed for 30 s. The solution was stirred additionally for 30 minutes. To improve adhesion force between PDMS and hydrogel, the PDMS-based ACFAL integrated with SFAT was treated with benzophenone (BZP) by first mixing 10% w/w BZP with acetone for 60s via vortexing followed by 60s of sonication to ensure complete incorporation of BZP in solvent. Subsequently, the solution was pipetted onto the surface of PDMS and exposed to air for solution to evaporate for 10 min. Upon complete evaporation, the PDMS surface was washed gently with DI water three times to remove excess BZP crystalline solids formed and dried with O_2_ air gun before depositing the bioadhesive hydrogel for curing. Lastly, bioadhesive hydrogel was integrated with SFAT-ACFAL by cross-linking the hydrogel solution under UV light for 15 minutes (∼4.21 J).

### Characterization of Bioadhesive hydrogel

#### Adhesion strength of bioadhesive hydrogel with skin and PDMS

The adhesion strength of the bioadhesive hydrogel was evaluated modified ASTM F2255-05 and ASTM F2256-05 methods through custom-developed and integrated testing machine (FB5, Torbal) with 90°-peeling off test. The samples were prepared with dimensions of 20 x 50 x 2 mm (width x length x thickness), and the backside of each sample was affixed with Kapton film (7413D, 3M) to prevent stretching during peeling. To measure the adhesion between the skin and bioadhesive hydrogel, the samples were gently attached onto a skin, and then peeled off at a 90° angle from the skin at a speed of 68 mm/min. To measure the adhesion between PDMS and bioadhesive hydrogel, PDMS was initially deposited and cured on a glass substrate mold (width: 50 mm, length: 76 mm). Then, a BZP treatment process was conducted. Using a similar 90°-peeling off test, the substrate was mounted and performed to compare adhesion force with and without BZP-treatment between the hydrogel and PDMS (**Fig. 3i**).

#### Weight Loss

To measure the dehydration characteristics of the hydrogel, a weight loss test was conducted. A circular-shaped bioadhesive hydrogel and a commercial gel (Aquasonic 100, Parker) sample were prepared (diameter: 19 mm, thickness: 1 mm). The weight of each was measured over time both in a typical room environment (∼41%, ∼23 °C) and inside a container with high humidity (∼65%, ∼23 °C) (**Supplementary Fig. 10**). The weight loss of the samples (W_l_) was calculated using the equation W_l_ (%) = (W_t_ – W_i_)/ W_i_ x 100, where W_i_ and W_t_ denote, respectively, the initial weight of the sample and the weight of the sample at different times.

#### Acoustic characteristics of hydrogel

Acoustic properties of the bioadhesive hydrogel were characterized by measuring and estimating the acoustic time-of-flight difference of ultrasound transmission through water, PET, and hydrogel between transducer and hydrophone. A single cycle sine wave pulse was generated using a 3-level beamformer transmitter circuit (TX7316, Texas Instrument) with a supplied driving voltage of ± 20 V. To measure the acoustic time-of-flight of the hydrogel, a 3-mm thick hydrogel was prepared with a mold consisting of PET film and Ecoflex frame (**Supplementary Fig. 11**) and measurements were performed over a period of 7 days. The purpose of the Ecoflex frame was to maintain the thickness of the hydrogel and to prevent the penetration of the water into the hydrogel when measuring in the water tank. Between measurements, the Ecoflex frame was removed temporarily, and the hydrogel samples were stored in a room environment (humidity: ∼30%, temperature: ∼23°C). Then, when measurements were taken again, the Ecoflex frame was placed around the hydrogel again to prevent water from entering the hydrogel. The acoustic time-of-flight was measured by placing the hydrogel samples between transducer and hydrophone in a water tank. The acoustic speed of the hydrogel was estimated by following equations^52^:

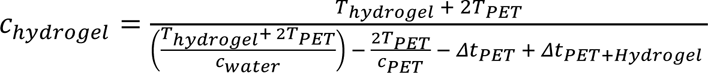

Where T_Hydrogel_ is the thickness of the bioadhesive hydrogel, T_PET_ is the thickness of the PET film, c_Water_ is the speed of sound in water (1500 m/s), c_PET_ is the speed of sound in PET film (polyethylene, high density: 2430 m/s, ^82^), ΔT_PET_ is ToF difference between with and without PET film, ΔT_PET+Hydrogel_ is ToF difference between with and without hydrogel samples.

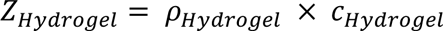

Where Z_Hydrogel_ is the acoustic impedance of the bioadhesive hydrogel, *ρ*_Hydrogel_ is the density of the hydrogel, c_Hydrogel_ is the speed of sound of the hydrogel.

#### Long-term acoustic stability

To assess the long-term acoustic stability of the bioadhesive hydrogel, the ultrasound intensity of the bioadhesive hydrogel integrated SFAT was measured over time. The number of circular-shaped hydrogels were prepared and stored in a container with high humidity (∼65%, ∼23°C). At specific time intervals, each bioadhesive hydrogels were taken out of the container and attached to a bare PZT transducer. All measurements were conducted under the deionized water. To prevent rapid swelling of the AMPS-based bioadhesive hydrogel upon contact with water, bioadhesive on the bare PZT transducer was covered with a thin Ecoflex cap (thickness: 0.5 mm). Then, the attenuation of ultrasound intensity due to the bioadhesive hydrogel over time was measured using a custom setup 3-axis hydrophone acoustic scanning system.

### Device Integration of MiniUlTra

Fabricated SFAT-ACFAL was connected via low temperature solder (NP510-LT HRL1, Kester) to a BNC cable and housed in a custom-designed 3D printed casing (PLA Galaxy, Prusa), which was lined with copper shielding (1181, 3M) and grounded to the BNC shielding layer for electromagnetic shielding purposes. To integrate SFAT-ACFAL with the bioadhesive hydrogel, The hydrogel solution was then poured to a thickness of 1 mm. Subsequently, the bioadhesive hydrogel on the SFAT-ACFAL was cross-linked under UV light for 15 minutes. Finally, the integrated device was completed by removing the mold (**Supplementary Fig. 8**).

### Characterization of tFUS on sensory-evoked potentials in S1 using SFAT-ACFAL

#### Participants

The Institutional Review Board (IRB) at University of Texas at Austin approved all experimental procedures under the study (STUDY00003279). Five healthy volunteers (4 male, 1 female, aged 24-36 with a mean age of 27.4 ± 5.1 years) provided written informed consent to participate in the study. Volunteers were screened for contraindications and neurological impairment and all subjects were right-hand dominant.

#### Experimental setup

Participants were positioned and seated in an adjustable height chair, where their right forearm is fully extended and supported in supination. Four 10-20 EEG electrode sites (C3, CP1, P3, CP5) were connected for recording somatosensory evoked potentials. During testing, subjects were initially stimulated by FES with varying currents (8-25 mA, 200 µs) to obtain the minimum threshold necessary to elicit muscle contraction of the right contralateral side. The SFAT-ACFAL was applied topically to CP3 manually with administration of ultrasound gel (Aquasonic 100, Parker) as interface to the scalp, which was then held in place using medical tape. Additionally, three electrical stimulation electrodes (2” Round, Reserv) were placed on the right contralateral arm (ground electrode on elbow, bipolar electrodes axially paired on the wrist via palpation of median nerve). The electrodes were connected to a functional electrical stimulation (FES) system (RehaMove3, Hasomed) for median nerve stimulation (MN), which was controlled externally by custom Python software.

tFUS treatment condition stimulation occurring 100 ms before MN stimuli (360µs ON and 640µs OFF, PRF 1kHz, Pulse Duration 500ms ON 500ms OFF) was controlled by programming of microcontroller (Uno, Arduino), which was connected to trigger the ultrasound generator (BBBoq, Image Guided Therapy System), FES system (MN stimuli), and EEG amplifier for time-locked epoch events during somatosensory evoked potentials (SEP) (**Supplementary Fig. 12**). Custom Python code was developed to integrate all systems together in addition to use of LabStreamingLayer (LSL) to stream and log EEG data into dataframe with external data including trigger and metadata. Subjects were then subjected to three blocks of trials, where each block consisted of four trials (FUS-/FES-, FUS-/FES+, FUS+/FES-, FUS+/FES+) and each trial lasted 3 mins. Within each trial, 30 s of baseline recording occurs before 120 s of sham/FUS followed by 30s of rest recording to ensure sufficient buffered data for post-recording cleaning. Total recording session time was approximately 1 h.

#### Electroencephalography

Subjects recruited were invited to a dedicated EEG recording room with minimal electronics for minimizing electromagnetic interferences. Tape ruler was used to measure the distance between nasion-to-inion and left-right preauricular points to determine electrode positioning according to the 10-20 system for EEG recording. Marker was used to indicate the position of C3, CP1, P3, CP5 for EEG and CP3 for tFUS targeting (**Fig. 4b**). Subsequently, rubbing alcohol was applied carefully at the sites before conductive hydrogel electrodes (H124SG, Kendall) were applied carefully to the scalp to ensure minimal obstruction of hair. Impedance per electrode was measured using commercial amplifier (eego MyLab, AntNeuro) to ensure it is less than 10kΩ. EEG data were digitized at 512 Hz and stored for offline analysis.

#### Statistical analysis of somatosensory evoked potentials

Digitized EEG data were analyzed offline by first filtering using a third-order butterworth bandpass filter (2-90 Hz) followed by a first-order butterworth bandstop filter (59-61 Hz) to remove DC offsets, mains interference, and high frequency noises. A total of 120 epochs per trial recorded was then extracted using custom MATLAB code using triggered signals as markers. Briefly, data were epoched around median nerve stimulus trigger, 200ms prior up to 500 ms after the trigger was extracted as a single epoch for analysis. Subsequently, the data was baseline corrected by subtracting the mean values from = −200 ms to 0 ms. For each epoch, inspection of artifacts using rejection criteria of absolute peak-to-peak amplitude threshold greater than 75 µV will be removed. Grand averaged epochs across 5 subjects and 15 trials were obtained to determine the effects of sham and FUS in SEP using SFAT-ACFAL elicited by MN stimuli. EEG biomarkers N20, P27, N33, P50, N70, P100, and N140 were extracted by obtaining the mean amplitude ± 2 ms the desired biomarker time event due to the difficulty to reliably identify SEP peaks accurately per trial. Statistical analyses were performed on mean peak-to-peak amplitudes for the N20/P27, N33/P27, P50/N33, N70/P50, P100/N70, N140/P100 and long potential (LP) components (**Supplementary Table 1-4**). These data were averaged across all trials and subjects and presented as mean ± s.e.m for different group conditions. Non-parametric statistical test using Wilcoxon signed-rank test was applied for SEP complexes to determine significance of treatment conditions.

#### Spatial-Temporal Analysis

Spatial-temporal frequency analysis was performed (MATLAB R2021a, The MathWorks) to decompose effects and changes in frequency spectrum due to S1 targeting using tFUS with SFAT-ACFAL as a function of time^83^. Short-time Fourier transform (STFT) was used with a window size of 4.8 ms and 2.3 ms overlap through Hamming window approach. Power of spectral data was then converted into power (dB). Comparison between treatment groups was performed by subtracting spectral epochs to observe dynamic changes in power with respect to frequency bands, where −3 dB and −6 dB corresponds to one-fold and two-fold decrease respectively (**Supplementary Fig. 6**).

### Long-term demonstration of tFUS neuromodulation of MiniUlTra

#### Participants

The Institutional Review Board (IRB) at University of Texas at Austin approved all experimental procedures under the study (STUDY00003279). Four healthy volunteers (4 male, aged 27-36 with a mean age of 32.5 ± 4.7 years) provided written informed consent to participate in the study. Volunteers were screened for contraindications and neurological impairment and all subjects were right-hand dominant.

#### Experimental setup

Subjects were invited to S1 targeted tFUS stimulation using SFAT-ACFAL for long-term study (Day 1, 7, and 28). For each session, subjects were pre-screened for contraindications before beginning the experiment. Four 10-20 EEG electrode sites (C3, CP1, P3, CP5) were connected for recording somatosensory evoked potentials (**Fig. 4b**). Minimum threshold for right contralateral hand movement due to MN stimulation was performed to obtain the minimum threshold necessary to elicit muscle contraction of the right contralateral side. The SFAT-ACFAL with bioadhesive hydrogel was applied to CP3 and held in place independently by its adhesive nature. For EEG recording stability, additional medical tape was used to fix EEG electrodes and transducers to prevent motion artifacts. Three electrical stimulation electrodes (2” Round, Reserv) were placed on the right contralateral arm similarly to the previous experiment for S1 targeting mentioned before. The electrodes were connected to a functional electrical stimulation (FES) system (RehaMove3, Hasomed) for MN stimulation.

tFUS treatment and sham conditions performed identically with the exception of subjects subjected to five blocks of trial, where each block consisted of two trials (FUS-/FES+; Sham, FUS+/FES+; Treatment) and each trial lasted 3 mins. Total recording session time was approximately 1 h.

#### Statistical analysis of somatosensory evoked potentials

A total of 120 epochs per trial recorded was then extracted using custom MATLAB code using triggered signals as markers. Grand averaged epochs across 4 subjects and 20 trials were obtained to determine the effects of sham and FUS in SEP using SFAT-ACFAL elicited by MN stimuli. EEG biomarkers N20 and P27 were extracted by obtaining the mean amplitude ± 2 ms the desired biomarker time event due to the difficulty to reliably identify SEP peaks accurately per trial. Statistical analyses were performed on mean peak-to-peak amplitudes for the N20/P27. These data were averaged across all trials and subjects and presented as mean ± s.e.m for different group conditions. Two-way ANOVA was applied for SEP complexes to determine significance of treatment conditions across days with treatment (hydrogel) and with control (commercial gel) in comparison to sham (no FUS) conditions within groups.

#### Study approval

All experiments were performed in compliance with the Institutional Review Board with approval at the University of Texas at Austin (STUDY00003279).

#### Code availability

The codes used for this study are available on GitHub at https://github.com/kevintang725/MiniUlTra-LSL.

## Acknowledgements

We thank Wynn Legon, Greg Fonzo, José del R. Millán, Samantha Santacruz and Yaoyao Jia for the discussions and advice. We would like to thank Nanshu Lu for her guidance and providing access to the use of the LPKF Protolaser U4 system. We would like to thank José del R. Millán for providing access to the RehaMove3 FES system. H.W. would like to acknowledge support from Alzheimer’s Association New to the Field (AARG-NTF) research grant, University of Texas at Austin Startup Fund, and NIH Maximizing Investigators’ Research Award (MIRA) (R35) grant. J.J would like to acknowledge support from the Human Frontier Science Program (HFSP) Fellowship. We acknowledge BioRender.com for the figures drawing.

## Author Contributions

Conceptualization: K.W.K.T., J.J., and H.W.; Methodology: K.W.K.T. and J.J.; Software: K.W.K.T.; Validation: K.W.K.T. and J.J.; Formal Analysis: K.W.K.T. and J.J.; Investigation: K.W.K.T., J.J., J-C.H., and M.M.Y.; Data Curation: K.W.K.T. and J.J.; Writing - Original Draft: K.W.K.T. and J.J.; Writing - Review & Editing: K.W.K.T., J.J., H.D., W.W., X.L., I.P., M.M.Y., W.H., W.D.M.B., A.R.L., B.A., and H.W.; Visualization: K.W.K.T. and J.J.; Project Administration: K.W.K.T and H.W.; Resources: N.L. and H.W.; Funding: H.W.

