## Supplementary Information for "Bioadhesive Hydrogel-Coupled and Miniaturized Ultrasound Transducer System for Long-Term, Wearable Neuromodulation"

### AUTHOR INFORMATION

Kai Wing Kevin Tang[1,†], Jinmo Jeong[1,†], Ju-Chun Hsieh[1,†], Mengmeng Yao[1], Hong Ding[1], Wenliang Wang[1], Xiangping Liu[1], Ilya Pyatnitskiy[1], Weilong He[1], William D. Moscoso-Barrera[1], Anakaren Romero Lozano[1], Brinkley Artman[1], Heeyong Huh[2], Preston S. Wilson[3], Huiliang Wang[1]*

### AFFILIATION

[1]*Department of Biomedical Engineering, Cockrell School of Engineering, The University of Texas at Austin, Austin, Texas 78712, United States.*

[2]*Department of Aerospace Engineering and Engineering Mechanics, Cockrell School of Engineering, The University of Texas at Austin, Austin, Texas 78712, United States.*

[3]*Walker Department of Mechanical Engineering, Cockrell School of Engineering, The University of Texas at Austin, Austin, Texas 78712, United States.*

[†]These authors contributed equally to this work.
*Corresponding author

Corresponding Author: Dr. Huiliang (Evan) Wang
Department of Biomedical Engineering, Cockrell School of Engineering, The University of Texas at Austin, Austin, Texas 787112, United States


Keywords: (ultrasound, neuromodulation, hydrogel, wearable devices)

***Experimental Section/Methods***

**PDMS Spin-coat Calibration.** Control of layer thickness when fabricating ACFAL for the SFAT was done by developing a calibration curve. Utilizing the same fabrication process, glass substrates were initially patterned with 36 um thick copper tape using laser etching (LPKF, U4 Laser) via transfer printing method (**Supplementary Fig. 3**). Glass substrates were cleaned and prepared by first submerging into a 1000mL beaker filled with acetone and sonicated in an ultrasonic sonicator to remove particulates for 5 min. Substrates were then removed, rinsed with distilled water and submerged in methanol for 5 min of sonication. The substrates were then rinsed with distilled water before blow dried with purified nitrogen gas. Substrate was spin-coated with a sacrificial layer (Omnicoat, Kayaku Advanced Materials) for 30s at 1000 RPM and 3 min of planarization before soft-baking at 200°C on a hotplate. Subsequently substrates were then spin-coated with 5 mL of PDMS (Sylgard 184); prepared by mixing 1:10 of curing agent with base elastomer and desiccated for 1 hour; at varying speeds (500-4000 RPM at 500 RPM increments) and cured on a hotplate at 90°C for 40 mins. Substrates were then placed in acetone filled beakers, for which it was sonicated for 5 min each to release the patterned PDMS mold. The PDMS mold was then reversely placed on a separate glass substrate for measurement. Each patterned mold substrate was then imaged (Axioscope 2 MAT, Zeiss) and measured using a profilometer (Dektak 150, Veeco) and layer thicknesses were determined accordingly (**Supplementary Fig. 7).**

**Ultrasound peak pressure calibration.** Measurement of peak pressure of the fabricated SFAT-ACFAL was performed by submerging the device into a degassed distilled water tank with hydrophone (Onda, Corporation, HGL-0200) connected to a preamplifier (Onda Corporation, AG-2010) with an amplification gain of 20dB. The hydrophone was aligned perpendicular to the face of the transducer. The water tank walls were lined with acoustic absorbing material (Precision Acoustic, Aptflex F28) to prevent acoustic reflections by absorption to reduce artifact and noise during measurement. Transducer was tested experimentally following sonication protocol used in the stimulation paradigm (340us ON and 640us OFF) at 650 kHz with varying amplitudes of 0-100% (**Figure 2e**). Measured voltage signals from hydrophone were connected and recorded by an oscilloscope (Siglent Technologies, SDS1202X-E), which was converted to pressure by calibration equation provided by the hydrophone manufacturer below.

$$Peak\ Pressure\ (Pa) = \frac{V_{measured}}{G(f)M_c(f)\frac{C_H}{C_H + C_C + C_A}}$$

$V_{measured}$: Voltage measured in oscilloscope via hydrophone and preamplifier

$G(f)$: Gain of preamplifier (Onda Corporation AG-2010, $G$ = 10 at 650kHz)

$M_c(f)$: End of cable nominal sensitivity of hydrophone (Onda Corporation HGL-0200, $M_c$ = 45nV/Pa at 650kHz)

$C_H$: Input capacitance of hydrophone (Onda Corporation HGL-0200, $C_H$ = 13pF at 650kHz)

$C_C$: Capacitance of right-angle connector between hydrophone and pre-amplifier ($C_C$ = 1.6 pF)

$C_A$: Input capacitance of pre-amplifier (Onda Corporation AGL-2010, $C_A$ = 6.3 pF at 650kHz)

Figure 14. Demonstration of the wearability of MiniUlTra. . ........ 12

**Table. 1. SEP Complexes in C3 channel.** Mean amplitudes of SEP complexes recorded from C3.

| **C3** | Mean Amplitude ± s.e.m. (μV) | | |
|---|---|---|---|
| SEP Complex | Sham | tFUS | P value |
| P27-N20 | 0.289 ± 0.083 | 0.086 ± 0.073 | 0.018* |
| N33-P27 | 0.154 ± 0.058 | 0.227 ± 0.073 | 0.241 |
| P50-N33 | 0.156 ± 0.083 | 0.016 ± 0.084 | 0.583 |
| N70-P50 | -0.571 ± 0.107 | -0.381 ± 0.114 | 0.583 |
| P100-N70 | 0.278 ± 0.182 | 0.075 ± 0.150 | 0.135 |
| N140-P100 | -0.009 ± 0.135 | 0.068 ± 0.164 | 0.714 |
| Late Potential | -0.260 ± 0.176 | -0.173 ± 0.185 | 0.295 |

**Table. 2. SEP Complexes in P3 channel.** Mean amplitudes of SEP complexes recorded from P3.

| **P3** | Mean Amplitude ± s.e.m. (μV) | | |
|---|---|---|---|
| SEP Complex | Sham | tFUS | P value |
| P27-N20 | 0.803 ± 0.221 | 0.469 ± 0.247 | 0.006** |
| N33-P27 | 0.662 ± 0.163 | 0.704 ± 0.141 | 0.413 |
| P50-N33 | 0.048 ± 0. 417 | 0.086 ± 0.495 | 0.951 |
| N70-P50 | -0.455 ± 0.199 | -0.327 ± 0.286 | 0.855 |
| P100-N70 | 0.185 ± 0.306 | 0.068 ± 0.227 | 0.583 |
| N140-P100 | -0.858 ± 0.281 | -0.526 ± 0.508 | 0.501 |
| Late Potential | -0.707 ± 0.429 | -0.567 ± 0.502 | 0.135 |

**Table. 3. SEP Complexes in CP1 channel.** Mean amplitudes of SEP complexes recorded from CP1.

| **CP1** | Mean Amplitude ± s.e.m. (μV) | | |
|---|---|---|---|
| SEP Complex | Sham | tFUS | P value |
| P27-N20 | 0.458 ± 0.109 | 0.266 ± 0.038 | 0.0637 |
| N33-P27 | 0.769 ± 0.131 | 0.814 ± 0. 140 | 0.426 |

| | | | |
|---|---|---|---|
| P50-N33 | -0.228 ± 0.261 | -0.344 ± 0.305 | 0.761 |
| N70-P50 | -0.693 ± 0.146 | -0.415 ± 0.301 | 0.760 |
| P100-N70 | -0.204 ± 0.269 | -0.187 ± 0.290 | 0.807 |
| N140-P100 | -0.201 ± 0.378 | -0.411 ± 0.463 | 0.357 |
| Late Potential | -0.600 ± 0.484 | -0.302 ± 0.513 | 0.267 |

**Table. 4. SEP Complexes in CP5 channel.** Mean amplitudes of SEP complexes recorded from CP5.

| **CP5** | Mean Amplitude ± s.e.m. (µV) | | |
|---|---|---|---|
| SEP Complex | Sham | tFUS | P value |
| P27-N20 | 0.222 ± 0.056 | 0.089 ± 0.068 | 0.020* |
| N33-P27 | 0.093 ± 0.036 | 0.153 ± 0.042 | 0.325 |
| P50-N33 | 0.156 ± 0.083 | 0.016 ± 0.084 | 0.067 |
| N70-P50 | -0.572 ± 0.107 | -0.382 ± 0.114 | 0.172 |
| P100-N70 | 0.278 ± 0.182 | 0.075 ± 0.150 | 0.057 |
| N140-P100 | -0.009 ± 0.135 | 0.069 ± 0.164 | 0.714 |
| Late Potential | -0.261 ± 0.176 | -0.173 ± 0.185 | 0.067 |

**Table. 5. ACFAL parameters.** Boundary radii dimensions for air-cavity Fresnel lens

| **$r_1$ (µm)** | **$r_2$ (µm)** | **$r_3$ (µm)** |
|---|---|---|
| 4940.47 | 7174.91 | 9011.83 |

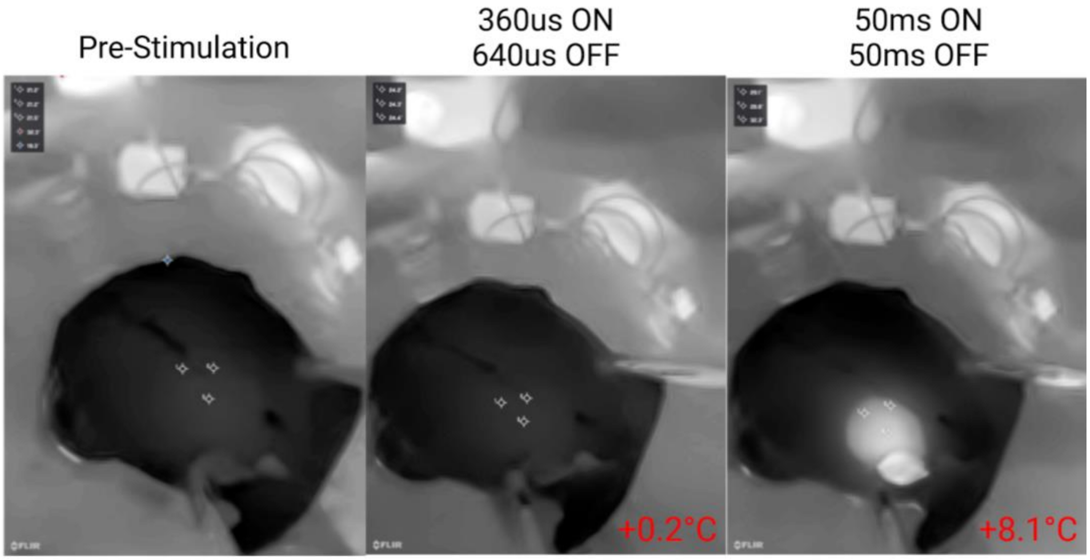

**Figure 1. Thermal heating effect of SFAT-ACFAL.** Infrared camera to measure thermal heating of MiniUlTra on macaque skull under varying stimulation conditions.

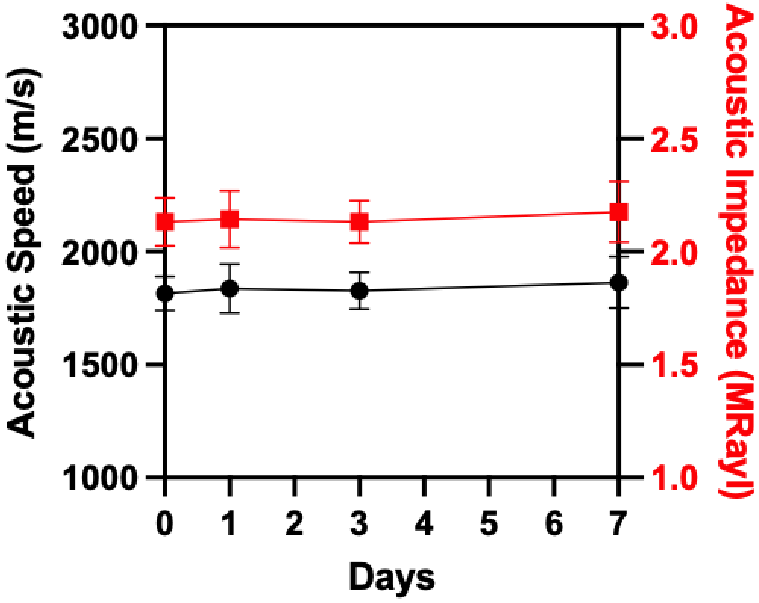

**Figure 2. Acoustic characterization of bioadhesive hydrogel.** Acoustic speed (black) and impedance (red) of the hydrogel for 7 days.

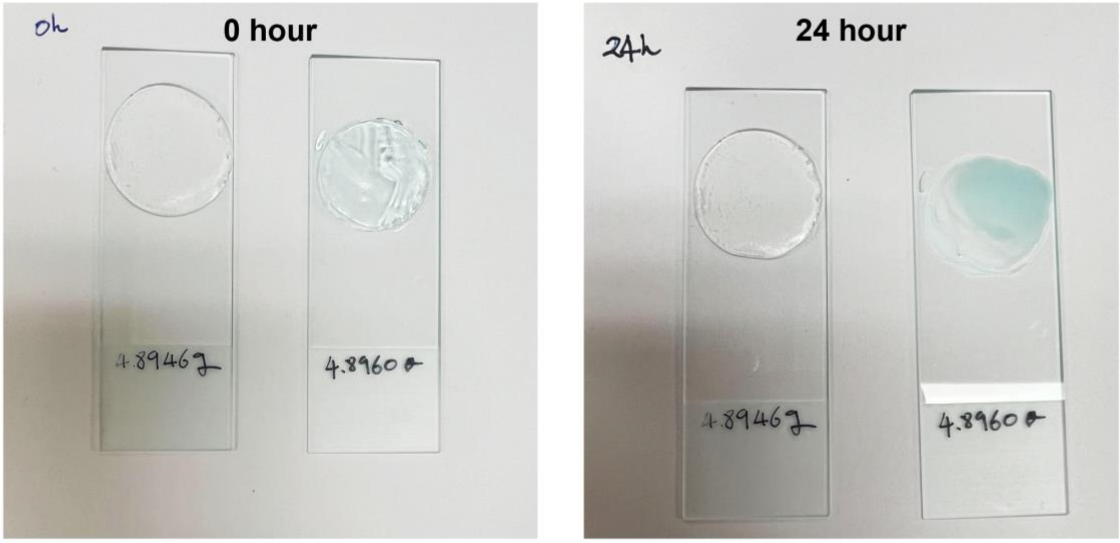

**Figure 3.** Comparison photographs of the dehydration state between the bioadhesive hydrogel (left) and commercial gel (right) over 24 hours.

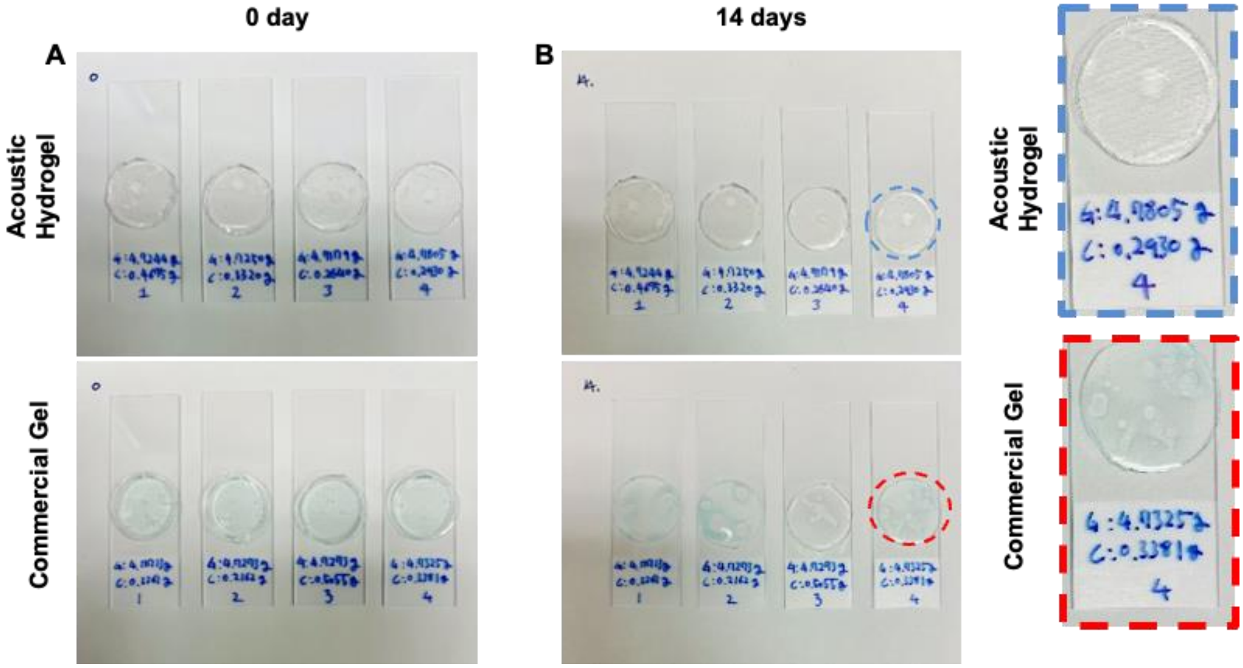

**Figure 4.** Comparison photographs of the dehydration state between (**A**) the bioadhesive hydrogel and (**B**) commercial gel over 14 days.

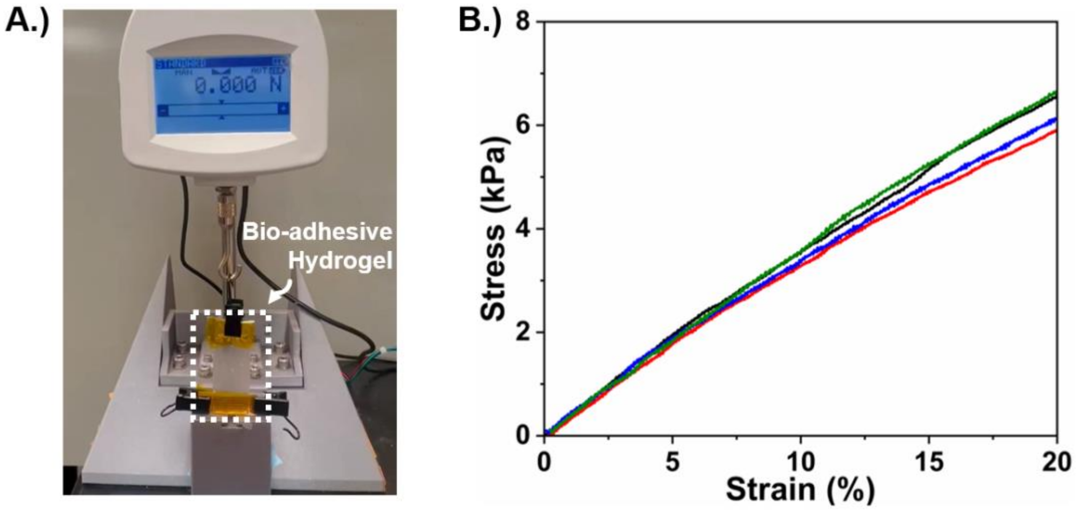

**Figure 5. Young's modulus of acoustic hydrogel A)** Experimental setup for measuring the adhesion force of the bioadhesive hydrogel. **B)** strain-stress curve of the bioadhesive hydrogel.

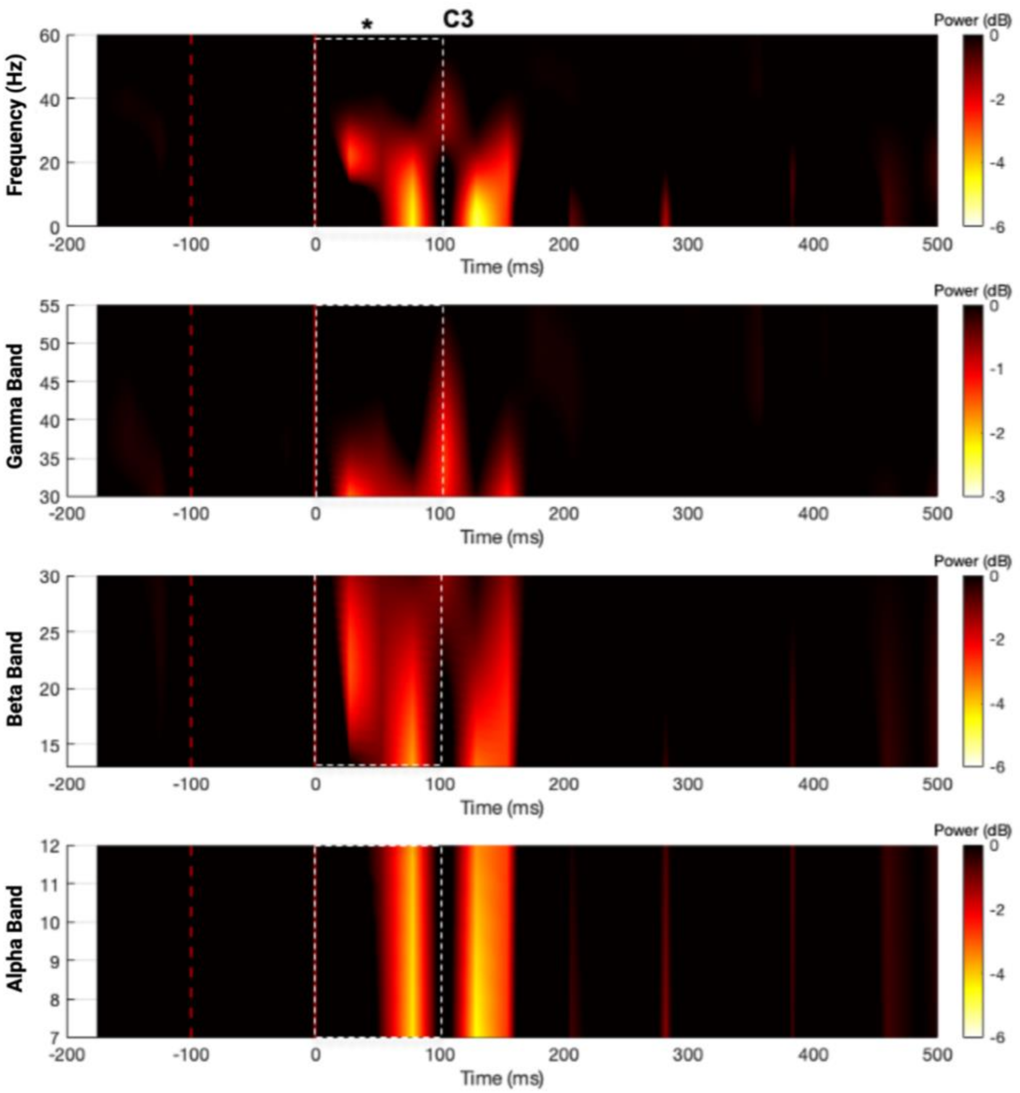

**Figure 6. Spatial-Temporal analysis of MiniUlTra in neuromodulation of sensory-evoked potential (SEP).** The application of MiniUlTra towards the S1 region during median nerve stimulation indicated a decreased power of alpha and beta band baseline activity recorded from EEG sites C3 within 100 ms onset from stimulus. Attenuation in the power of short-latency evoked gamma-band activity occurred also within 70 ms compared to sham (outlined in white dashed line). Attenuation was calculated by subtracting FUS from sham group's spatial-temporal spectrum of grand-averaged epochs and taking the logarithmic power in decibels (-3 dB and -6 dB corresponds to 50% and 25% power of maximum respectively).

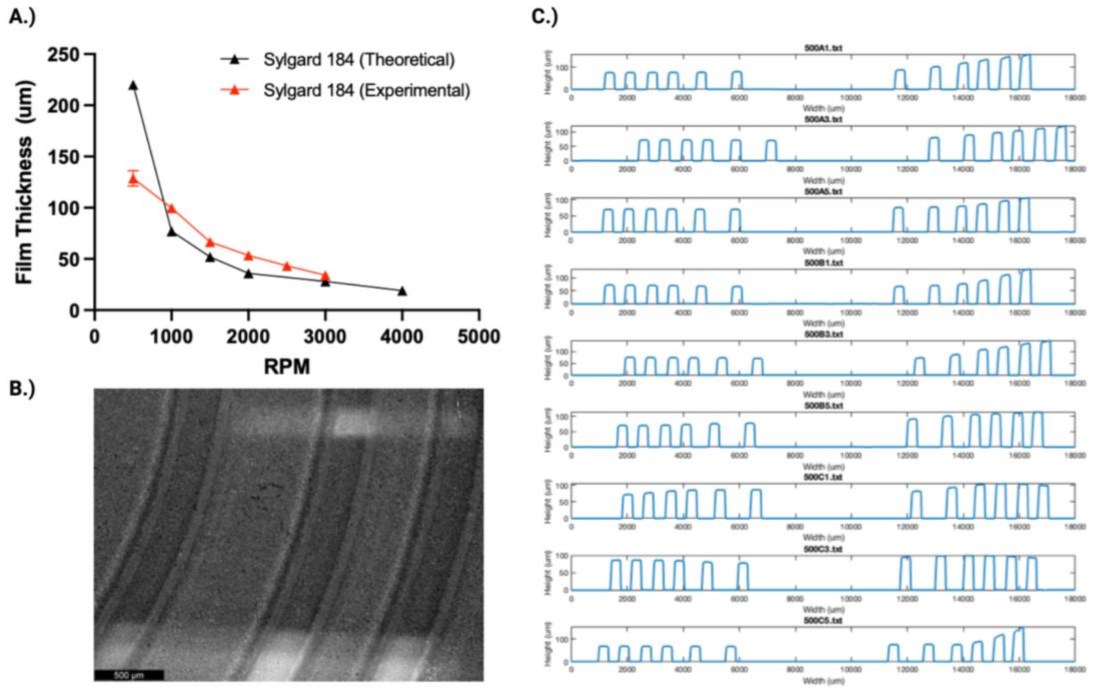

**Figure 7. PDMS Spin-coating Calibration. A)** PDMS spin-coating calibration curve. B**)** Microscope imaging of patterned ACFAL. C**)** Profilometer measurements of samples.

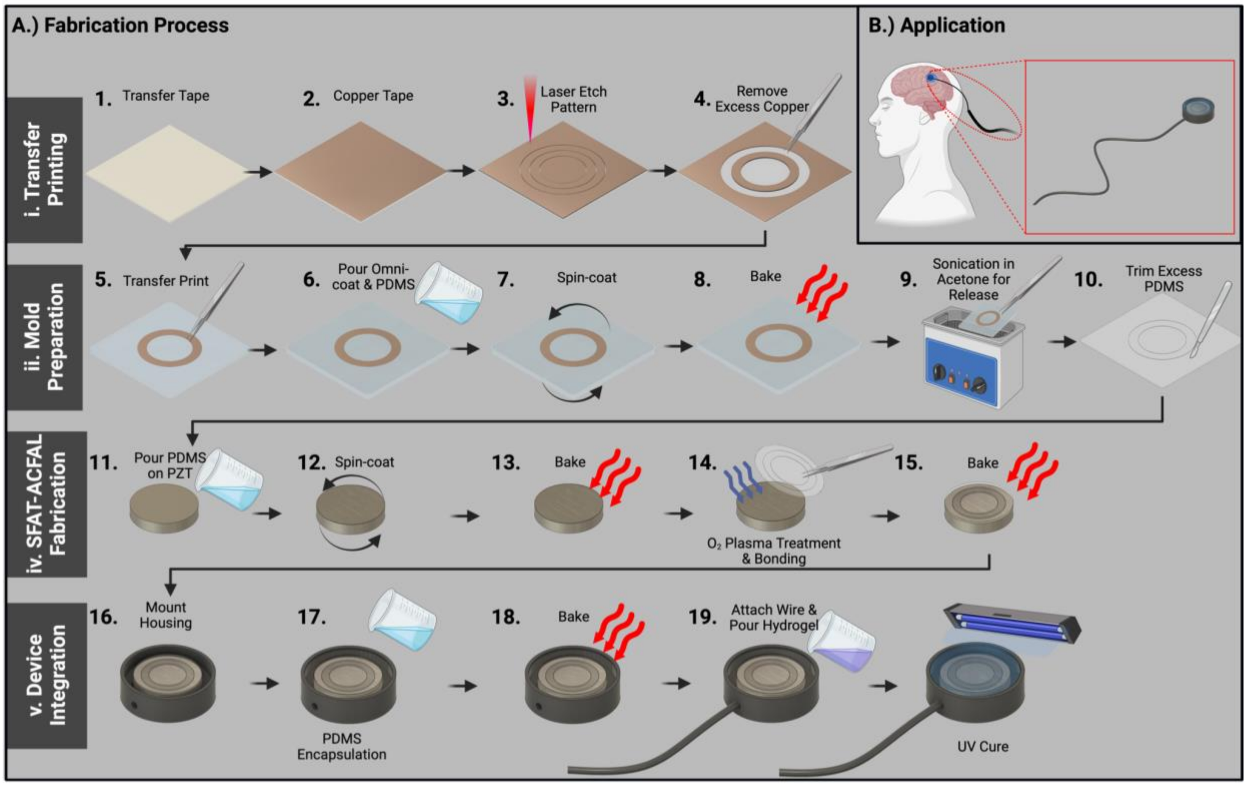

**Figure 8. Fabrication Procedure of MiniUlTra A)** Novel fabrication process of device without the need of traditional lithography. **i)** Transfer printing of laser etched copper tape pattern onto glass substrate to develop desired negative pattern. **ii)** Development of ACFAL using transfer printed mold with PDMS via sacrificial layer of Omnicoat. **iii)** Development of SFAT by $O_2$ plasma treatment of ACFAL for reverse bonding. **iv)** Device integration of SFAT-ACFAL into housing and connectors encapsulated with PDMS and bioadhesive hydrogel coupling. **B)** Application of MiniUlTra towards somatosensory cortex (S1).

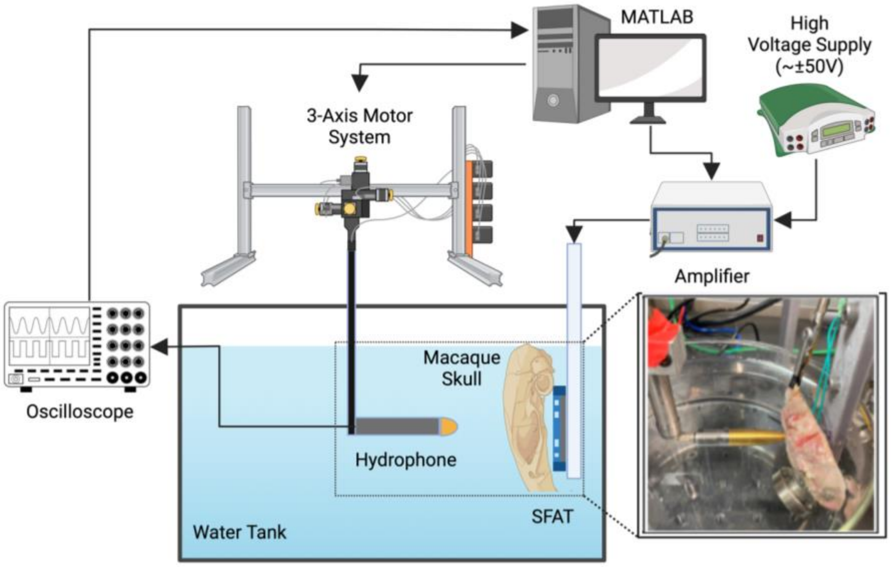

**Figure 9. Acoustic Field Measurement.** Experimental setup for measuring and characterizing SFAT-ACFAL with and without macaque skull.

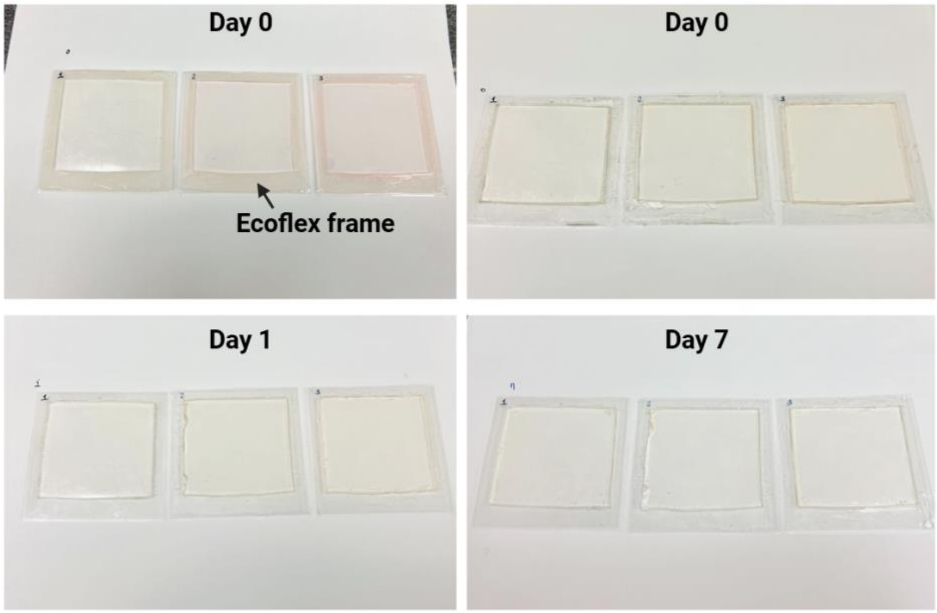

**Figure 10.** Photographs of the hydrogel samples for acoustic speed measurement for 7 days. The acoustic speed in hydrogel was measured with an Ecoflex frame. Between measurements, the Ecoflex frame was removed temporarily, and the hydrogel samples were stored in a room environment (humidity: ~30%, temperature: ~23°C)

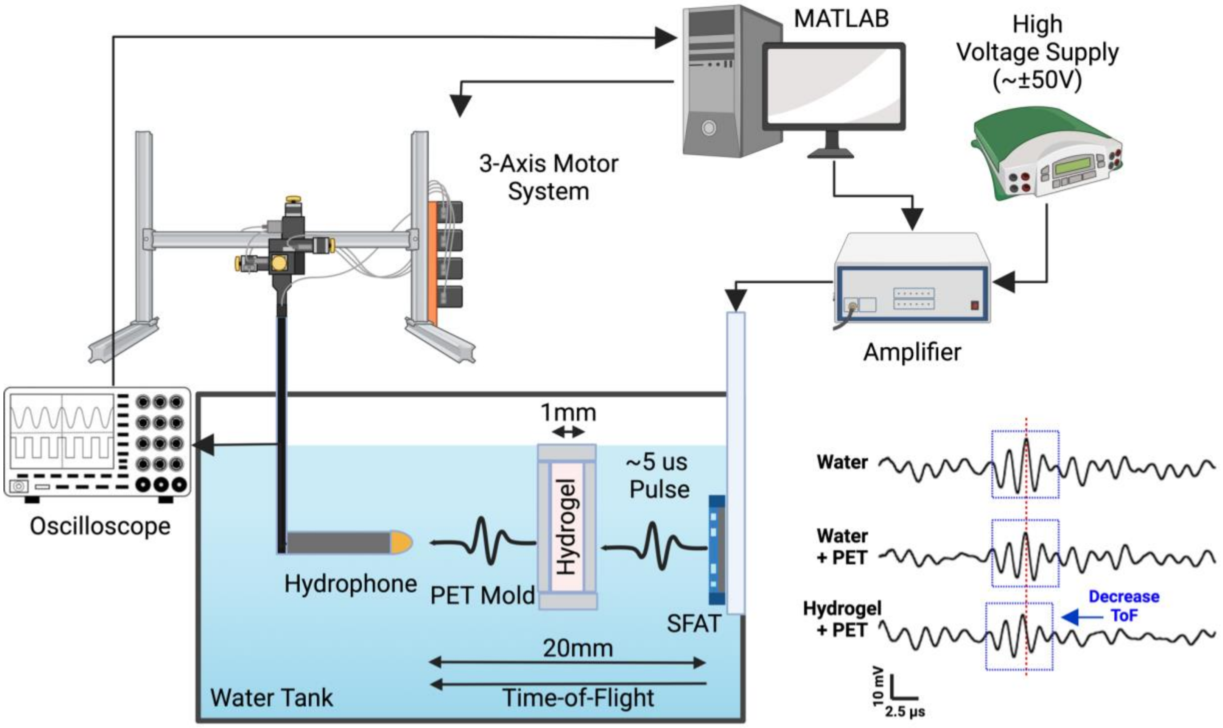

**Figure 11.** Experimental setup for estimating the acoustic speed of the bioadhesive acoustic hydrogel

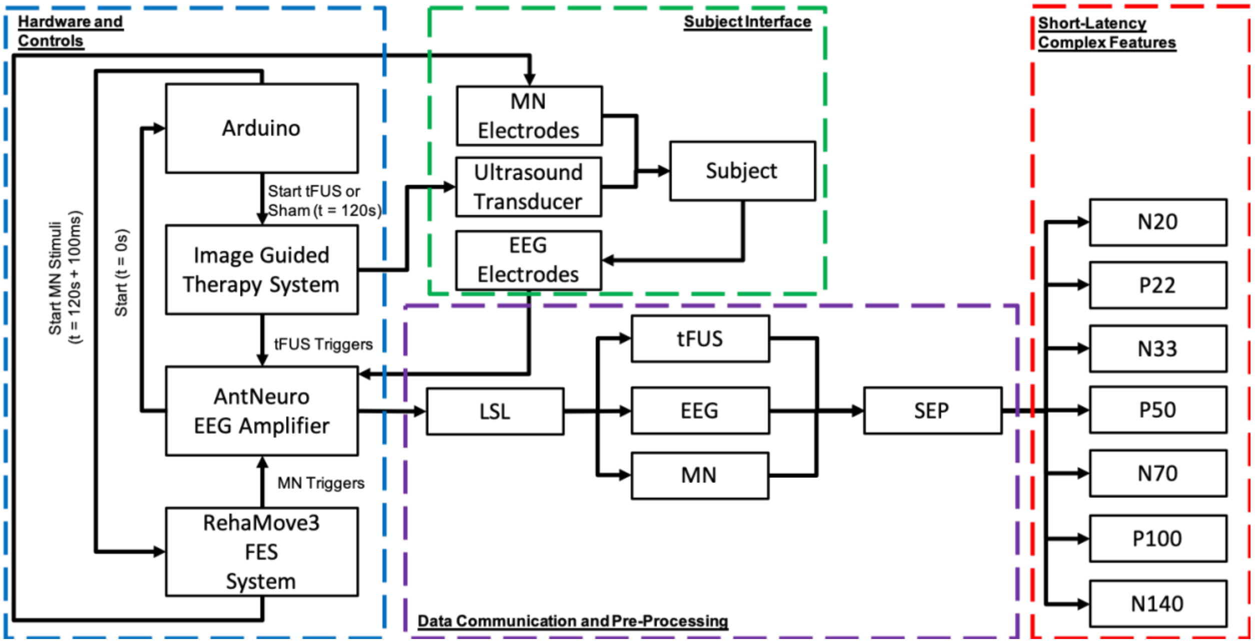

**Figure 12. Somatosensory Evoked Potential Experimental Setup.** Arduino Uno was programmed with a stimulation paradigm and triggers ultrasound (BBBoq, Image Guided Therapy System), median nerve stimulation (RehaMove3 FES System) and EEG recording triggers (AntNeuro). Four channels (C3, CP1, P3, CP5) of EEG data were recorded and saved via LabStreamingLayer. Epochs were extracted to evaluate SEPs and short-latency complex features were obtained.

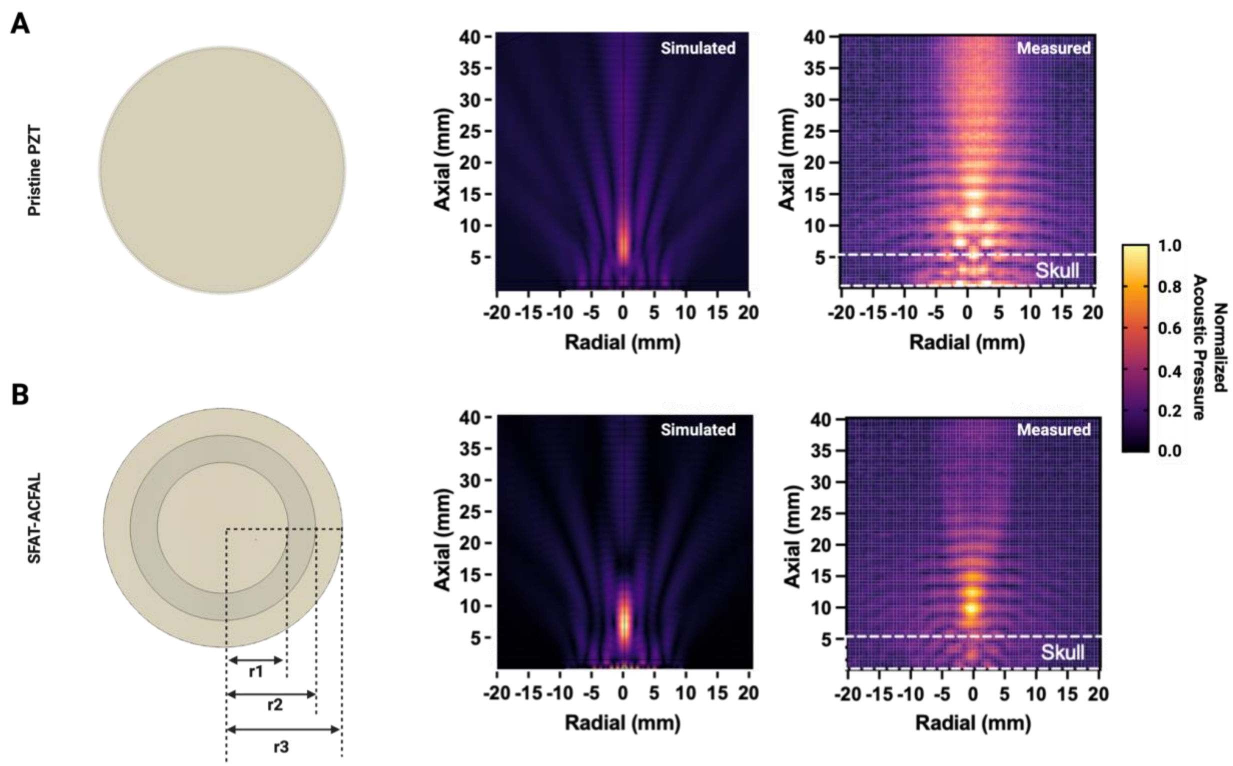

**Figure 13. Design and simulation of SFAT-ACFAL. A)** Comparison of simulated and measured results with pristine DL-47 PZT. Focal spot is more dispersed and lower intensity based on simulation, where measured result is highly scattered and beam profile shows no focality. **B)** Comparison of designed SFAT-ACFAL on DL-47 PZT. Simulated and measured results show comparable similarities, with increased acoustic intensity and higher spatial resolution of focal spot at focal depth of 10 mm.
